# Interictal spikes during spatial working memory carry helpful or distracting representations of space and have opposing impacts on performance

**DOI:** 10.1101/2024.11.13.623481

**Authors:** Justin D. Yi, Maryam Pasdarnavab, Laura Kueck, Gergely Tarcsay, Laura A. Ewell

**Author notes:** These authors contributed equally (Justin D. Yi, Maryam Pasdarnavab). Senior author (Laura A. Ewell).

## Abstract

In temporal lobe epilepsy, interictal spikes (IS) – hypersynchronous bursts of network activity – occur at high rates in between seizures. We sought to understand the influence of IS on working memory by recording hippocampal local field potentials from epileptic mice while they performed a delayed alternation task. We found that IS disrupted performance when they were spatially non-restricted and occurred during running. In contrast, when IS were clustered at reward locations, animals performed well. A machine learning decoding approach revealed that IS at reward sites were larger than IS elsewhere on the maze, and could be classified as occurring at specific reward locations – suggesting they carry informative content for the memory task. Finally, a spiking model revealed that spatially clustered IS preserved hippocampal replay, while spatially dispersed IS disrupted replay by causing over-generalization. Together, these results show that IS can have opposing outcomes on memory.

## INTRODUCTION

Temporal lobe epilepsy (TLE) is the most common focal epilepsy syndrome, and is often comorbid with cognitive impairments. Deficits in episodic memory and working memory are common^1–6^. One clearly pathological feature of memory processing in TLE is that memory tasks promote interictal spikes (IS)^7^, hyper-synchronous network events observed as large spikes in local field potential (LFP) recordings. The recruitment of elevated IS rates during memory tasks suggests that the network mechanisms that promote IS may be hijacking dynamics that are typically engaged by memory processes^8^. Whether or not the underlying neural dynamics of IS resemble healthy processing and are thus helpful for memory performance, or are divergent enough to constitute interfering signals is an open question.

The exact timing of IS during a memory task has different impacts on performance^9–11 12^, but the mechanisms of such timing selective impairments are poorly understood. It is known that spatial working memory task performance is disrupted in rodent models of TLE^13^, however, it is not known if the exact timing of IS with respect to different task phases contributes to the memory deficit. One possibility is that different phases of memory tasks rely on neural dynamics that are more or less similar to IS dynamics, and therefore more or less susceptible to interference. For example, as animals navigate the maze, the hippocampal network is in a low synchrony state and engages in movement related theta oscillations which orchestrate the sequential activation of individual neurons over second long time scales^14–18^. On the other hand, as animals consume reward or sit quietly during delay phases of the task, the network shifts to a high synchrony state and engages in brief periods of physiological network synchrony called sharp wave ripples (SWR)^19^, which replay memory relevant ensembles of neurons at compressed time scales^20^. It is possible that if IS recruit hippocampal neurons in a similar enough manner to how SWR do, and if they are activated in the same task contexts as SWR, then they may play a similar role in memory.

There is evidence to suggest that there are parallels between IS and SWR neuronal dynamics and that, in principle, they could partially fulfill analogous functional roles. For example, both recruit ensembles of CA1 pyramidal cells, are accompanied by brief fast oscillations measured in cell layer LFP (pathological high frequency oscillations for IS and ripple oscillations for SWR), and coordinate with the cortex via sleep spindles^21–24^. Furthermore, in epilepsy, when IS rates go up during a memory task, SWR rates go down^25^, indicating that IS may actually replace SWR. If IS were simply hijacking SWR during working memory, it is plausible that task processes that rely on such activity would be maintained in epilepsy. Of course, this would depend on whether IS recruit proper memory relevant ensembles, which is not always the case^26^. A further complication is that SWR play diverse roles in memory tasks, such as amplifying salient cues, remote replay of past animal positions and rewards, or preplay of future trajectories^27–32^. Thus it is possible that IS can mimic some, but not all SWR dynamics.

During active locomotion, IS may interact with theta oscillations in a way that SWR do not. During theta oscillations in healthy animals, co-active ensembles of neurons are relatively small, due to rhythmic inhibition, which ensures precise encoding of environments^33,34^. Notably, SWR, which recruit larger co-active ensembles, are very rarely observed during locomotion-related theta states^19^ due to the elevated levels of inhibition which suppress synchrony and neuromodulatory signals that are strong during theta^35^. This suggests that hyper-synchrony during locomotive theta states is not a normal feature of healthy hippocampal circuits. Interestingly, when inhibition is reduced experimentally, theta oscillations during running can grow into large amplitude spikes that resemble bursts of interictal spikes^36^. Furthermore, in epilepsy, IS have been reported to encroach on theta states^23^, however whether IS during theta states impact memory is unknown. In other words, IS may create interference by promoting aberrant population level synchrony during theta states.

Given the relationship between IS and SWR, the differing roles of SWR in memory, and the observations that IS can encroach on theta states, there are several possible ways in which IS could mechanistically alter spatial working memory. To study these questions, we employed *in vivo* electrophysiology in freely moving TLE mice while they performed the delayed alternation spatial working memory task. We characterized when IS occur with respect to task phase to interpret impacts on behavioral performance. To gain further mechanistic understanding of our observed results, we developed a behavioral model to explain interactions between IS rate and task engagement. Furthermore, we employed a machine learning-based decoding approach to study whether IS features might be informative of task demands. Finally, a spiking neural network model was created to test the impact of IS during working memory on hippocampal replay.

## RESULTS

### Epileptic mice exhibit persistent focal interictal activity

To test how hippocampal dynamics during spatial working memory are impacted by interictal activity, saline (control) or kainic acid (KA) injected mice were implanted with drivable microelectrodes which were positioned in the hippocampus over several days (Table 1). Once electrodes were in their final positions mice were video monitored to determine rates of seizures and interictal discharges during restful periods (total of 13.0 ± 2.0 monitoring hours/mouse). As expected, mice injected with KA experienced frequent subclinical seizures (12 ± 11 of seizures/hour, Table 2), confirming that they suffer from focal temporal lobe epilepsy (Figure 1A). In addition to subclinical seizures, we observed seemingly sustained interictal spiking that was categorized into two types: solitary interictal spikes (IS) and chains of spikes called Brief Interictal Rhythmic Discharges (BIRDs). Events were classified as solitary IS or BIRDs based on inter-spike intervals similar to^37^ (Figure 1B-D, Table 3).

**Table 1:** Interictal and ictal spike detection parameters for each animal. Settings were optimized according to the procedure detailed in the Methods.

| Animal ID | Type | Threshold (μV) | Low pass-band (Hz) | High pass-band (Hz) | F<sub>1/2</sub> Score | Precision (%) | Recall (%) |
| --- | --- | --- | --- | --- | --- | --- | --- |
| m1 | Tetrode | 900 | 2 | 400 | 0.45 | 50% | 32% |
| m2 | Tetrode | 900 | 16 | 400 | 0.92 | 97% | 76% |
| m3 | Tetrode | 1900 | 1 | 400 | 0.86 | 90% | 71% |
| m4 | Tetrode | 1300 | 1 | 400 | 0.58 | 69% | 35% |
| m5 | Tetrode | 1100 | 1 | 400 | 0.99 | 99% | 99% |
| m6 | Tetrode | 700 | 1 | 400 | 0.78 | 83% | 62% |
| m7 | Probe | 1700 | 1 | 400 | 0.70 | 75% | 56% |

**Table 2:** Rates of spontaneous seizures during monitoring. Mean, maximum, and minimum seizure rates were pooled across all monitoring sessions. Seizures were defined as trains of spikes with inter-spike intervals less than 2 s with a train duration of at least 10 s. *Animal m7 was monitored during the light-cycle, whereas all others were monitored during the dark-cycle.

| Animal ID | Number of Monitoring Sessions | Mean Seizure Rate (hr<sup>-1</sup>) | Max. Rate (hr<sup>-1</sup>) | Min. Rate (hr<sup>-1</sup>) |
| --- | --- | --- | --- | --- |
| m1 | 10 | 6.5 | 18.0 | 0.0 |
| m2 | 12 | 5.0 | 15.0 | 0.0 |
| m3 | 12 | 10.9 | 21.7 | 3.7 |
| m4 | 14 | 7.2 | 29.1 | 0.0 |
| m5 | 15 | 7.4 | 26.8 | 1.0 |
| m6 | 15 | 11.5 | 24.7 | 1.0 |
| m7* | 14 | 36.9 | 50.1 | 6.0 |

**Table 3:** Rates of interictal events during monitoring sessions. *Animal m7 was monitored during the light-cycle, whereas all others were monitored during the dark-cycle.

| Animal ID | Number of Monitoring Sessions | Mean Interictal Spike Rate (Hz) | Std. Dev. Interictal Spike Rate (Hz) | Mean Solitary Spike Rate (Hz) | Std. Dev. Solitary Spike Rate (Hz) | Mean BIRD Rate (Hz) | Std. Dev. BIRD Rate (Hz) |
| --- | --- | --- | --- | --- | --- | --- | --- |
| m1 | 10 | 0.792 | 0.589 | 0.034 | 0.013 | 0.041 | 0.019 |
| m2 | 12 | 0.227 | 0.139 | 0.020 | 0.006 | 0.030 | 0.012 |
| m3 | 12 | 0.468 | 0.136 | 0.049 | 0.005 | 0.072 | 0.008 |
| m4 | 14 | 0.356 | 0.203 | 0.039 | 0.006 | 0.064 | 0.031 |
| m5 | 15 | 0.214 | 0.139 | 0.018 | 0.006 | 0.030 | 0.013 |
| m6 | 15 | 0.266 | 0.067 | 0.033 | 0.009 | 0.043 | 0.008 |
| m7* | 14 | 0.708 | 0.136 | 0.038 | 0.007 | 0.069 | 0.008 |

**Figure 1:**
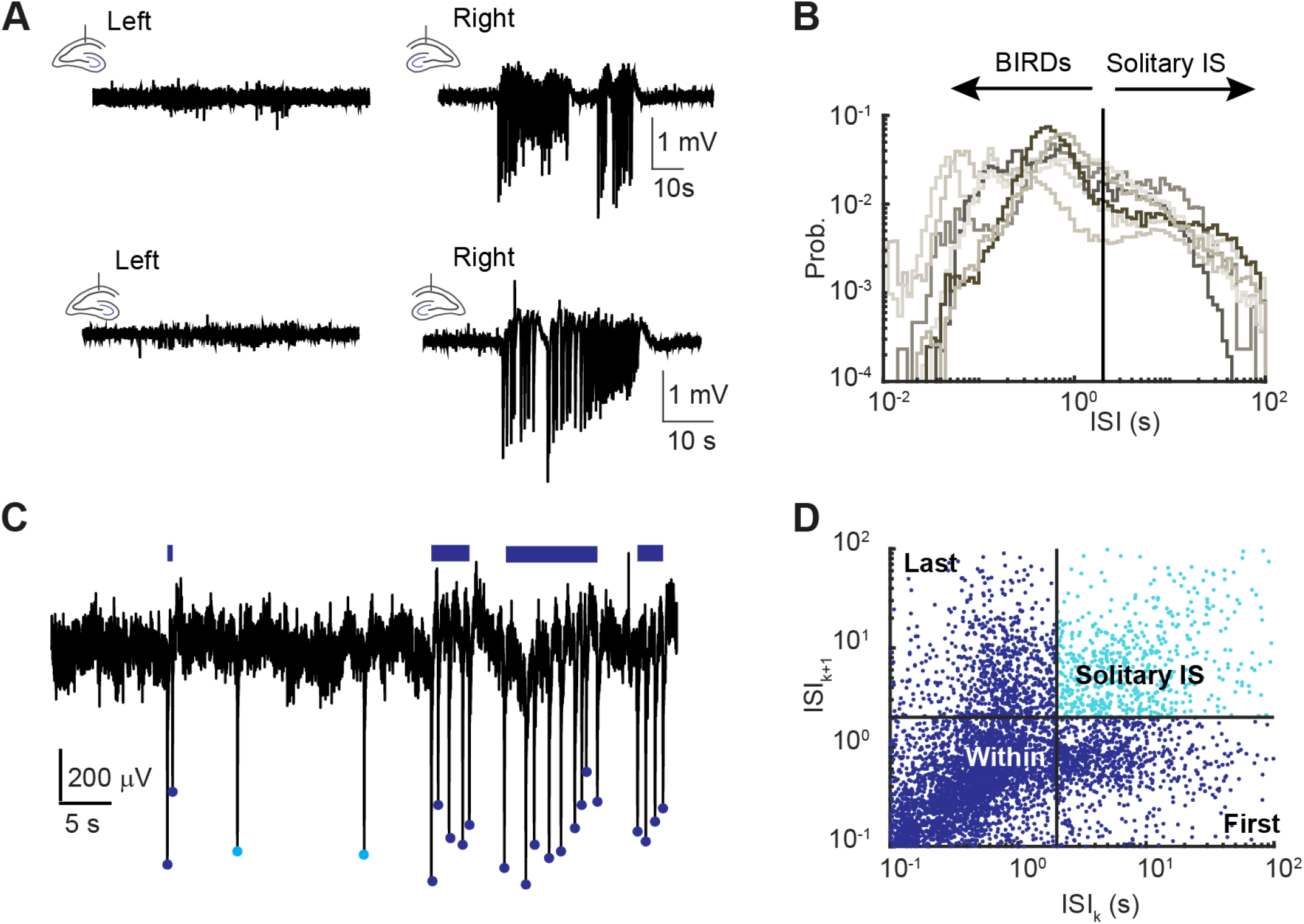
KA mice exhibit spontaneous seizures and interictal activity. (A) Two examples of seizures recorded from the hippocampus bilaterally where the right hemisphere was injected with KA. Spikes were detected continuously both during subclinical seizures and in the interictal period. (B) For each KA animal, the inter-spike intervals (ISI) between each interictal spike (IS) was used to classify IS as solitary IS or chains of IS called brief rhythmic interictal discharges (BIRDs). IS with ISI greater than 2 seconds were considered solitary, and less than 2 seconds as part of BIRDs. (C) Examples of solitary IS (light blue dots) and BIRDs (dark blue dots), with BIRD durations shown as bars. (D) A Poincaré plot shows a sampling of ISI pairs which can be divided into “First”, “Within” and “Last” spikes of BIRDs or solitary spikes using the same 2 second threshold as in (B).

### Epileptic mice have impaired performance on a spatial working memory task

In addition to video-LFP-monitoring, mice were recorded during daily behavior sessions comprising a spatial working memory task flanked by rest sessions. While performing the delayed alternation spatial working memory task, mice had to alternate between visiting two sides of a Figure-8 shaped maze (Figure 2A) to receive food (or liquid sucrose for m7) rewards with a 30 second delay period between trials (see methods for training details)^38^. Over the five sessions of testing, control mice (n=6) performed significantly better than KA mice (n=7) (repeated measures ANOVA, F(1,11)=7.25, p=0.021) (Figure 2B). The difference in behavior was also observed when averaging performance across the five sessions of testing (Figure 2C) (control, n = 6, 76.4 ± 3.1 %; KA, n = 7, 59.34 ± 5.2 %, unpaired t-test, d.f.= 11, t-stat = 2.69, p = 0.021). Notably, the KA group did not perform better than the chance level of 50% correct choices (one sample t-test, t-stat = 0.63, d.f.=6, p = 0.55), whereas control mice did perform significantly higher than chance level (one sample t-test, t-stat = 3.2, d.f.=5, p = 0.023) (Figure 2C). We also noted that despite poor overall performance, the KA group exhibited individual sessions of good performance (Figure 2D), suggesting that the mechanisms underlying poor performance may be dynamic.

**Figure 2:**
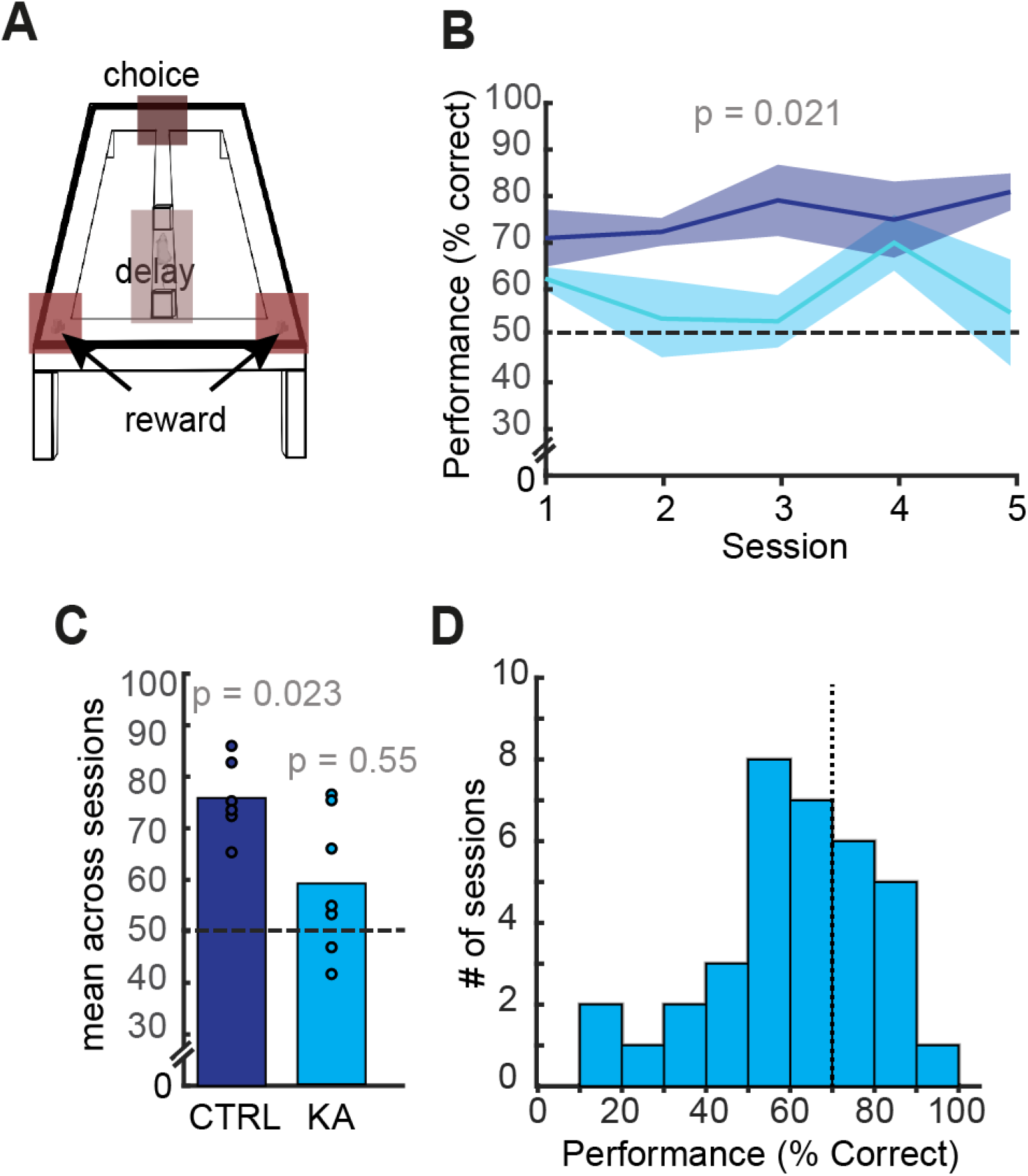
KA mice have impaired working memory performance. (A) The Figure-8 maze used for delayed spatial alternation with salient locations highlighted. (B) Control animals’ performance in the delayed alternation task (dark blue line) was significantly higher than that of KA animals (light blue). (C) Furthermore, the mean performance across all 5 sessions was higher than chance (50%) only for control (CTRL) animals. (D) Day-to-day performance of KA animals was variable but interspersed with “good” sessions (*>*70% performance dashed line).

### IS occur during working memory and their spatial distribution correlates with memory performance

To determine what mechanisms underlie impaired and variable memory performance in KA animals, we recorded hippocampal local field potentials during task performance. Animals had high rates of IS while performing the working memory task (0.50 ± 0.07 Hz, n = 35 sessions = 7 animals × 5 sessions) (Figure 3). Spikes either occurred as solitary interictal spikes (IS) (0.027 ± 0.003 Hz) or in BIRDs (0.035 ± 0.003 Hz). BIRDs were typically short in duration (4.5 ± 0.42 seconds) and comprised several spikes (14 ± 2 spikes). We noted that for some mice the spatial distribution of spikes were confined to specific areas of the maze, and were even consistent across sessions of memory testing (e.g., m1 and m7 3A). Other mice exhibited patterns of spiking that extended across large portions of the maze and were more variable session to session (e.g., m3 and m6). Consistent with this observation, we found that the spatial information of interictal spikes, which is a measure of how well spiking activity predicts mouse location, was quite variable across sessions (Figure 3B, left), with some sessions exceeding values of 2 bits/spike. Such high values of spatial information match those reported for individual place cells in healthy hippocampus^39^. High spatial information of interictal spikes was weakly, but significantly associated with a better performance on the working memory task (Figure 3B, right, p = 0.049; see Table 4 for further statistical details).

**Table 4:** GLM coefficients comparing spatial information (SI) to animals’ per session performance, adjusted *R*^2^ = 0.085, d.f. = 33, dispersion = 0.035.

| Parameter | Estimate | SE | tStat | p-value |
| --- | --- | --- | --- | --- |
| Intercept | 0.5 | 0.05 | 10.0 | $< 1 \times 10^{-5}$ |
| SI | 0.08 | 0.04 | 2.0 | 0.049 |

**Figure 3:**
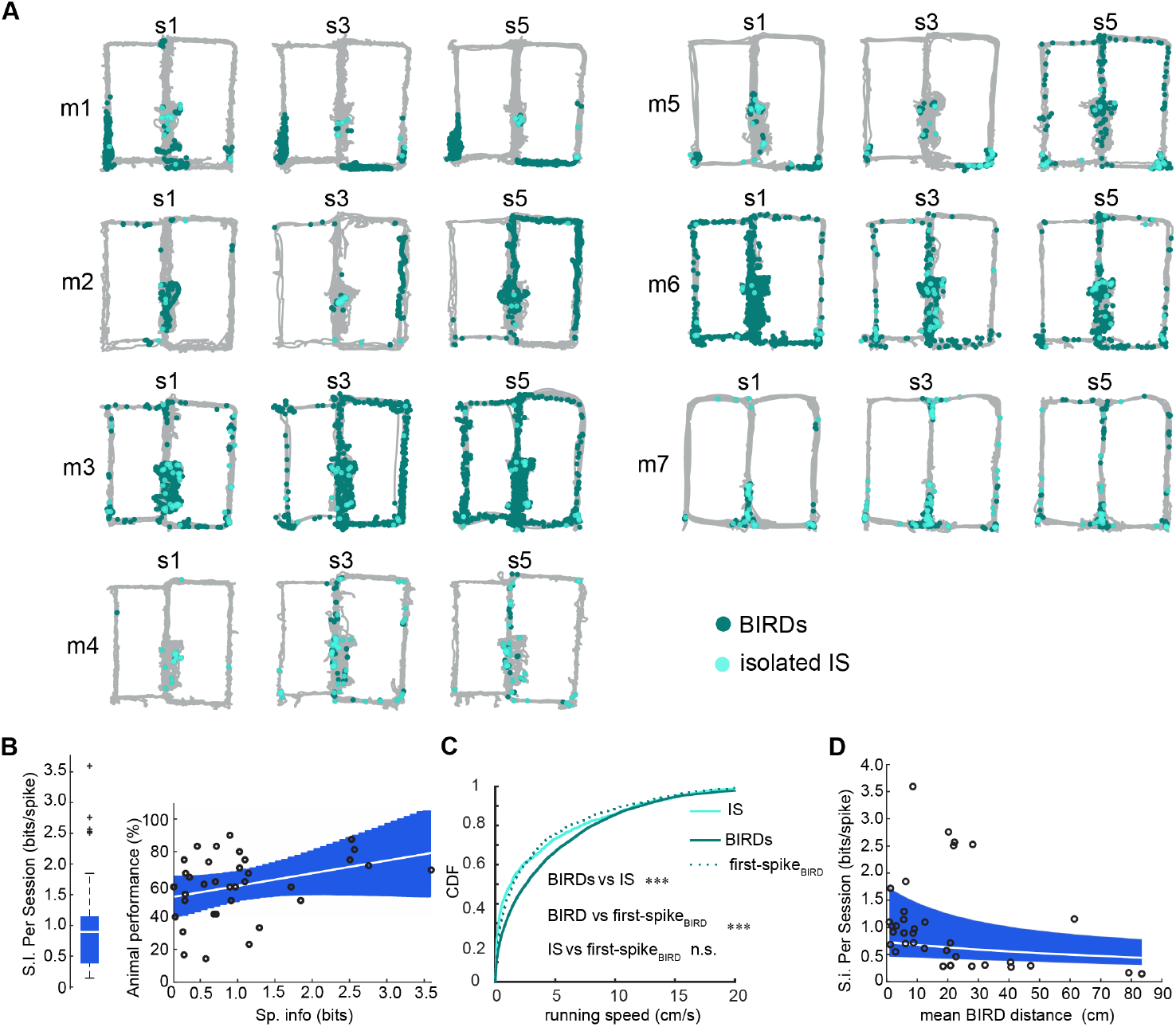
IS and BIRDs occur during execution of a working memory task. (A) The locations of BIRDs (dark blue dots) and solitary IS (light blue dots) on the maze for all animals on three of the five sessions. (B) The spatial information per interictal spike was computed for each session (N=35 = 7 mice × 5 sessions, left). Higher values of spatial information corresponded to a higher alternation performance predicted by a GLM (right) (p = 0.049, see Table 4 for details). (C) The running speed during solitary spikes (light blue), BIRDs (dark blue), and the first spike of each BIRD (dashed) was compared to reveal that BIRDs occur at faster running speeds than solitary IS or the first spike in each BIRD. Table 5 contains statistics for the comparisons shown in C, *** p*<*0.001, n.s. p*>*0.05. (D) Using a GLME, it was found that working memory sessions that had BIRDs associated with long running trajectories significantly explained lower values of spatial information (fixed-effect for distance term, p-value = 0.02, see Table 6). The marginal (unconditional) fixed effect mean and 95% CI are shown in the blue shaded region.

To further investigate contributions to the variable nature of spatial information of interictal spiking, we calculated running speeds at the times of IS and BIRDs. Solitary IS occurred during periods of rest as reported by others^22^, while BIRDs tended to occur when the mouse was running at faster speeds (Figure 3C, see Table 5 for statistics). Interestingly the first spike in a BIRD had a speed-tuning distribution that overlapped with solitary IS (Table 5), indicating that BIRDs may initiate from quiet restful states but can encroach onto running states if the animal begins movement mid-BIRD. We reasoned that BIRDs during running would drive lower spatial information, and indeed sessions with BIRDs that spanned larger distances on the maze were associated with lower total information per spike (Figure 3D, GLME fixed-effect for distance term, p-value = 0.02, Table 6 for more statistics).

**Table 5:** Running speeds of IS and BIRDs. Results of Wilcoxon rank sum test for the running speed distributions in Figure 3.

| Groups | p-value | z-value | Rank sum |
| --- | --- | --- | --- |
| IS vs BIRDs | $1 \times 10^{-16}$ | -8.3 | $6.6 \times 10^6$ |
| BIRD vs first spike of BIRD | $9 \times 10^{-15}$ | 7.8 | $1.3 \times 10^8$ |
| IS vs first spike of BIRD | 0.07 | -1.8 | $1.0 \times 10^6$ |

**Table 6:**
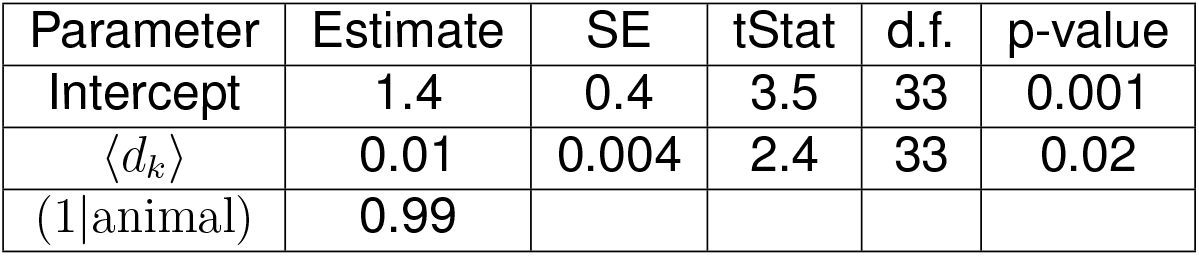
Coefficient values for the gamma GLME in Eq. 6, estimated dispersion was 0.08. Adjusted *R*^2^ = 0.82 SE = standard error, d.f. = degrees of freedom.

### The distribution of IS in the behavioral maze is augmented in specific spatial zones of the maze

To see whether IS were more likely to occur at specific maze locations, we divided the maze into “Delay,” “Choice,” “Reward,” and “Outer Arm” zones and calculated both the total time each animal occupied that zone and the IS rate in that zone (Figure 4A-B). The occupancy distribution was significantly different from the distribution of spikes in each zone (*χ*^2^ test, p-value = 2.6 × 10^−9^, dof = 1,*χ*^2^ stat: 35.44 Figure 4B), indicating that the interictal spike-generating process is non-stationary. To understand the zone-specific effects on the IS rate, we modeled the nonstationarity as a non-homogeneous Poisson process in which a “baseline” spike rate, *ρ*_*a*_, which is specific to each animal, is scaled by zone-specific gain factor, *η*_*z*_, unique to each zone but shared between all animals (Figure 4C).

**Figure 4:**
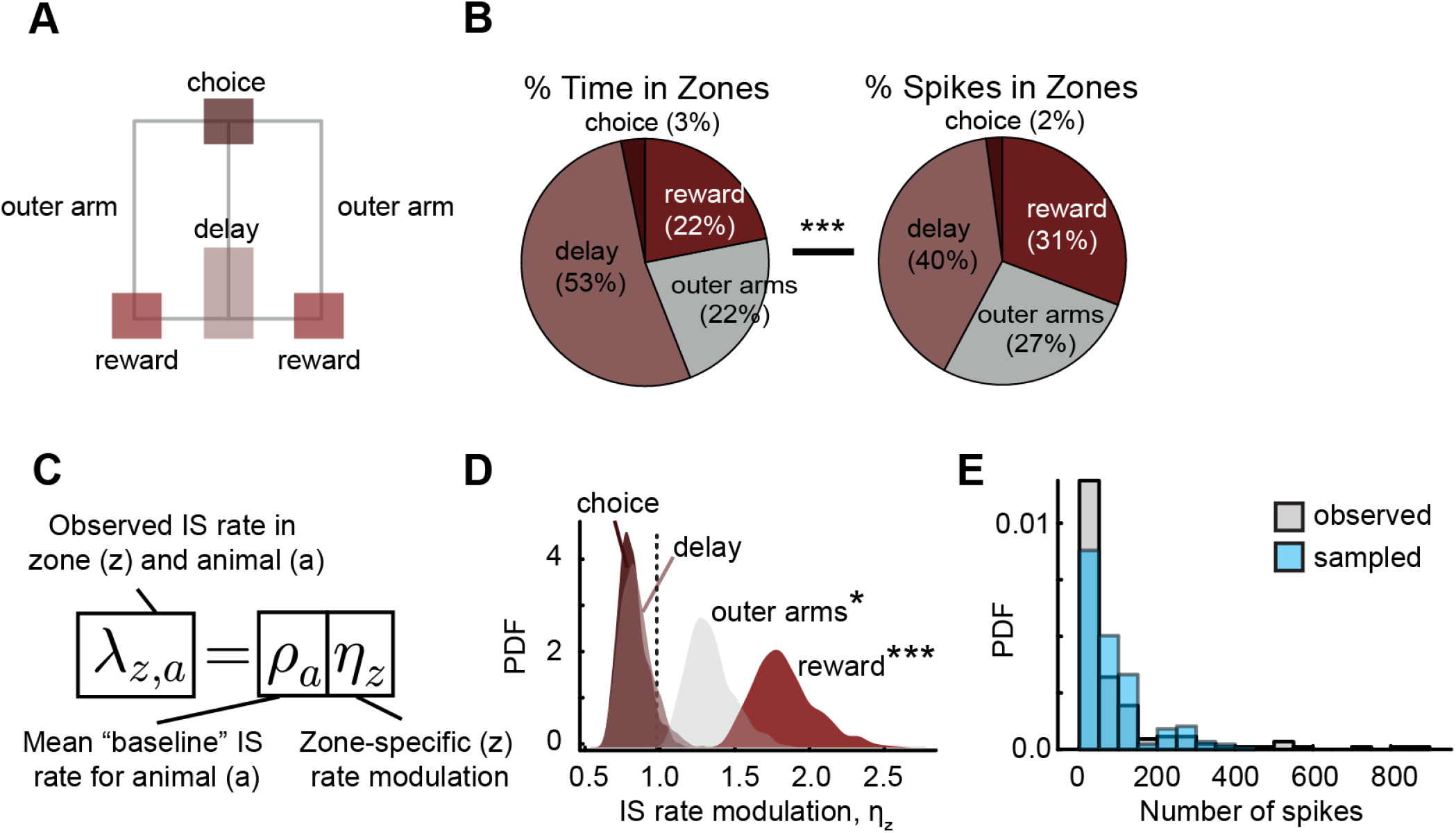
IS and BIRD rates are augmented in certain spatial zones of the maze. (A) For choice, delay, outer arm, and reward zones, (B) the proportion time spent and proportion of IS in each zone differed significantly from each other (*χ*^2^ test, p-value = 2.6 × 10^−9^, dof = 1, *χ*^2^ stat: 35.44). (C) To examine how each zone affected the IS rate of each animal, a Bayesian model was estimated (see Methods for details), where a zone-specific gain *η*_*z*_ = 1 was interpreted as a “neutral” effect. (D) The “outer arms” and “reward” zones had 95% highest posterior density (HPD) intervals of *η*_outer arm_ = [1.1, 1.6] and *η*_reward_ = [1.5, 2.3], respectively. (E) As a posterior predictive check, the distributions of IS spike counts actually observed were compared to those predicted by the Bayesian model. The bulk of the distributions (i.e. for means *<*400 spikes) agree whereas observed over-dispersion in the tails was not fully captured.

The posterior distributions of *η*_*z*_ in each zone were compared to a null value of 1 indicating the absence of a zone-specific modulatory effect on the IS rate. The “Reward” zone’s gain significantly deviated from 1 (Figure 4D; *η*_reward_ = [1.5, 2.3], 1 − *α* = 95% highest posterior density (HPD) interval, *N* = 7 animals × 5 sessions × 4 zones) and the “Outer Arm” zones’ term also deviated from 1 (*η*_outer arm_ = [1.1, 1.6] HPD interval). In other words, the IS rate was significantly elevated from baseline when the animal occupied reward zones and when the animal ran down outer arms to the reward zones, but the IS rate was consistent with baseline at all remaining locations on the maze. These results are consistent with our findings regarding spatial information, as we would expect that sessions with IS augmented at reward sites would have high spatial information, whereas sessions that had IS while the animal ran down outer arms of the maze would drive lower spatial information.

The model’s fit and inferences were inspected to assess model plausibility. We validated the model’s inferences by confirming that the distribution of posterior means of *ρ*_*a*_ (0.46 ± 0.23 Hz, n = 7 mice, mean ± 95% CI) agreed with the “naïve” time averaged IS rate (0.50 ± 0.07 Hz, n = 7 mice × 5 sessions), which was not explicitly given as data to the model. The mean values predicted by the model were compared directly to the observed data, where it was found the model distribution qualitatively agreed with the observed data (Figure 4E).

### Reward zone LFP discriminability predicts animals’ working memory performance

Given the significantly elevated IS rate in reward zones (Fig. 4D), which in some animals exhibited place cell like precision across sessions (Figure 3), we hypothesized that the IS LFP at reward zones may contain latent information regarding the location of the animal on the maze. Several studies have shown that features of the hippocampal LFP signal can be decoded to reveal a continuum of generating mechanisms^40–44^, and even into variables describing the animal’s behavioral state including position^45–47^ and social context^48^. After non-linearly embedding each IS LFP into a 2-dimensional space (Fig. 5A), a bagged ensemble of trees binary classifier^49^, sometimes referred to as a “random forest”^50^, was trained to discriminate between IS which occurred at reward sites versus those that did not. The classifier’s performance as measured by the receiver operating characteristic (ROC, see Supplemental Figure S1) area-under-the-curve (AUC), was able to predict the animal’s mean performance on the alternation task (Fig. 5B). Furthermore, when considering a classifier on only spikes that occurred in the reward zones, east and west reward sites could also be discriminated above chance level (Fig. 5C). This suggests that mice which generate IS in reward zones that are sufficiently distinct from IS in other locations on the maze have better spatial working memory, and that reward-IS carry spatial signals that are helpful for solving the task. This is consistent with reports that SWR in healthy animals recruit cells which encode locations near rewarded locations^27,31^. The effect of classifier AUC on predicting animals’ performance was consistent when controlling for mean spatial information, suggesting that both discriminability of reward-related IS and spatial information of IS are important and explain different aspects of the variance (Supplemental Table S1). Interestingly, IS which occurred in reward zones had significantly larger relative amplitudes than those that occurred in other locations on the maze (Fig. 5D). Similarly, when considering only IS within reward zones, the relative amplitudes for those which occurred during correct choices were also significantly larger than those during incorrect choices (Fig. 5D). This is consistent with reports that SWR in healthy animals at reward sites are larger in amplitude and longer in duration than at unrewarded locations^27^. Thus, the reward-related changes in IS features we have observed mirror those of reward SWR, suggesting that the decodability and amplitude differences we observe in IS may be driven by similar mechanisms that also recruit larger SWR and engage ensembles that encode locations near rewards.

**Figure 5:**
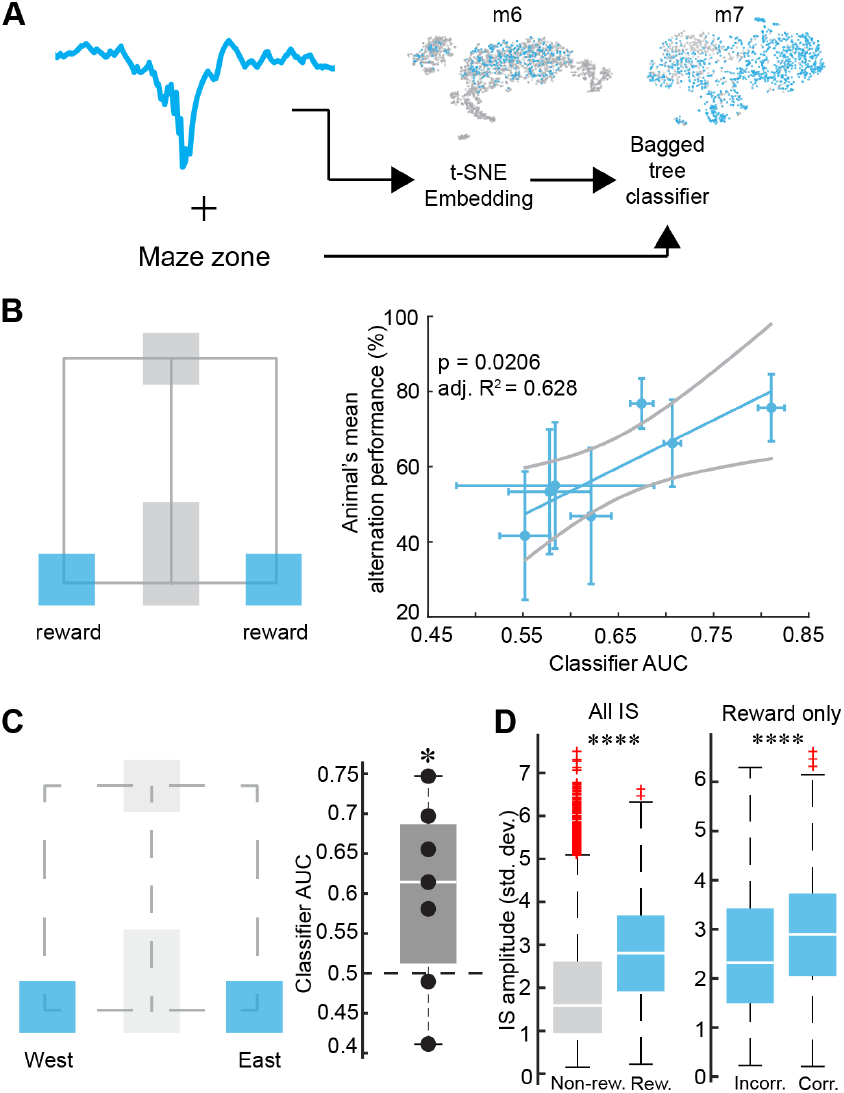
The ability to decode reward zone occupancy from the IS LFP predicts animals’ memory performance. (A) The normalized LFP from each IS was non-linearly embedded into a 2-dimensional space using the t-SNE algorithm^96^. A bagged ensemble of trees binary classifier^49,50^ was trained on the embedded IS LFP to decode whether the IS happened within a reward zone (blue) or not (gray dots). The t-SNE embedding of two representative animals’ LFP are shown. (B) The classification was evaluated using the receiver operating characteristic (ROC) area-under-the-curve (AUC). The AUC value was associated with the animal’s mean performance across the five sessions of behavior (p = 0.0206, see Table 7 for further details). Solid line is the mean and light lines are 95% CI of the regression model, error bars show 95% CI of individual data points. (C) The analysis was repeated for IS generated in reward zones only to see if animals maintained a representation of east v.s. west reward zones. The population mean of the AUC values was significantly greater than 0.5 chance level (tStat = 2.24, df = 6, sd = 0.12, p = 0.033, one-sided t-test; * p *<* 0.05). (D) The root-mean-squared amplitude, normalized by the standard deviation per given animal, was computed for each IS waveform. The left panel shows IS at reward zones was larger than all other zones (tStat = -40, df = 16299, sd = 1.3, two-sided t-test; **** p *<* 0.001). The right shows for spikes in reward zones only, IS during which the animal was rewarded were slightly larger (tStat = -8.2, df = 4721, sd = 1.2, p =*<* 0.001, two-sided t-test).

### Task-engagement state is related to performance and IS rates during the delay phase

A key phase of working memory is the delay phase. In our case, this corresponds to the 30 second period between trials when animals must maintain representations of the past to inform future decisions or ‘hold on’ to a future plan. In healthy animals, it is known that SWR during delay phases often replay locations of recently visited reward locations^32^, which is thought to support future decisions to not revisit that location on the next trial. Furthermore, interrupting SWR in between components comprising a multi-step task selectively impairs memory performance^29^, suggesting that SWR are critical for memory processes which take place on similar timescales as behavior. We therefore were interested in IS in the delay phase and whether or not delay phase IS play the role of delay phase SWR.

**Table 7:**
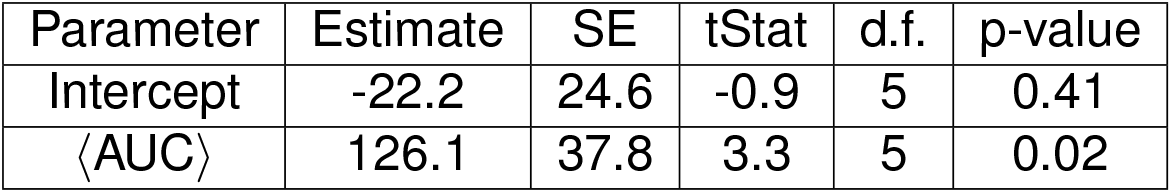
Coefficient values for the regression model in Figure 5B. Adjusted *R*^2^ = 0.628 SE = standard error, d.f. = degrees of freedom. Comparison against constant model: F statistic = 11.1, p = 0.0206

| Parameter | Estimate | SE | tStat | d.f. | p-value |
| --- | --- | --- | --- | --- | --- |
| Intercept | -22.2 | 24.6 | -0.9 | 5 | 0.41 |
| $\langle \text{AUC} \rangle$ | 126.1 | 37.8 | 3.3 | 5 | 0.02 |

First, we accounted for variations in engagement with the memory task which may co-vary with IS rates. Task engagement is known to fluctuate in healthy animals between distinct states with different error rates^51^. Therefore, we first estimated distinct task engagement states. Using the mice’s trial-to-trial performance, we inferred three discrete task-related behavioral states corresponding to low (p(Correct) =19%), medium (53%), and high (75%) success rates using a hidden Markov model scheme (HMM, Fig. 6). Naturally, the medium level state is consistent with a random guess, and the high-level engaged state corresponds to performing the task correctly with few errors. The low performance state is consistent with the strategy of perseveration, i.e. choosing the last visited site repeatedly. Within a single day, the mice typically transitioned from an initial “guessing” state to an “engaged” state, or relatively less often a “perseveration” state characterized by many errors in a row (Figure 6A). Control animals’ performance also was represented with an HMM (Supp. Fig. S2) of similar structure to that in Fig. 6B. Importantly, the probability of remaining in a perseveration state was lower in controls than epileptic mice (Supp. Fig. S3). In agreement with computational and psychophysical investigations of reaction time and decision certainty^52–54^, mean time to exit the delay zone was inversely related to the probability of correct choice as summarized in Table 8.

**Table 8:** The mean delay exit time (time to exit delay zone after 30 s interval elapsed) was estimated from 1000 samples drawn using the hierarchical bootstrap method^97^ for each discrete state estimated by the Viterbi algorithm.

| State (% corr.) | Mean delay exit time (s, 95% Conf. interval) |
| --- | --- |
| Perseveration (19%) | [11.9, 12.2] |
| Guess (53%) | [7.8, 7.9] |
| Engaged (75%) | [7.55, 7.6] |

**Figure 6:**
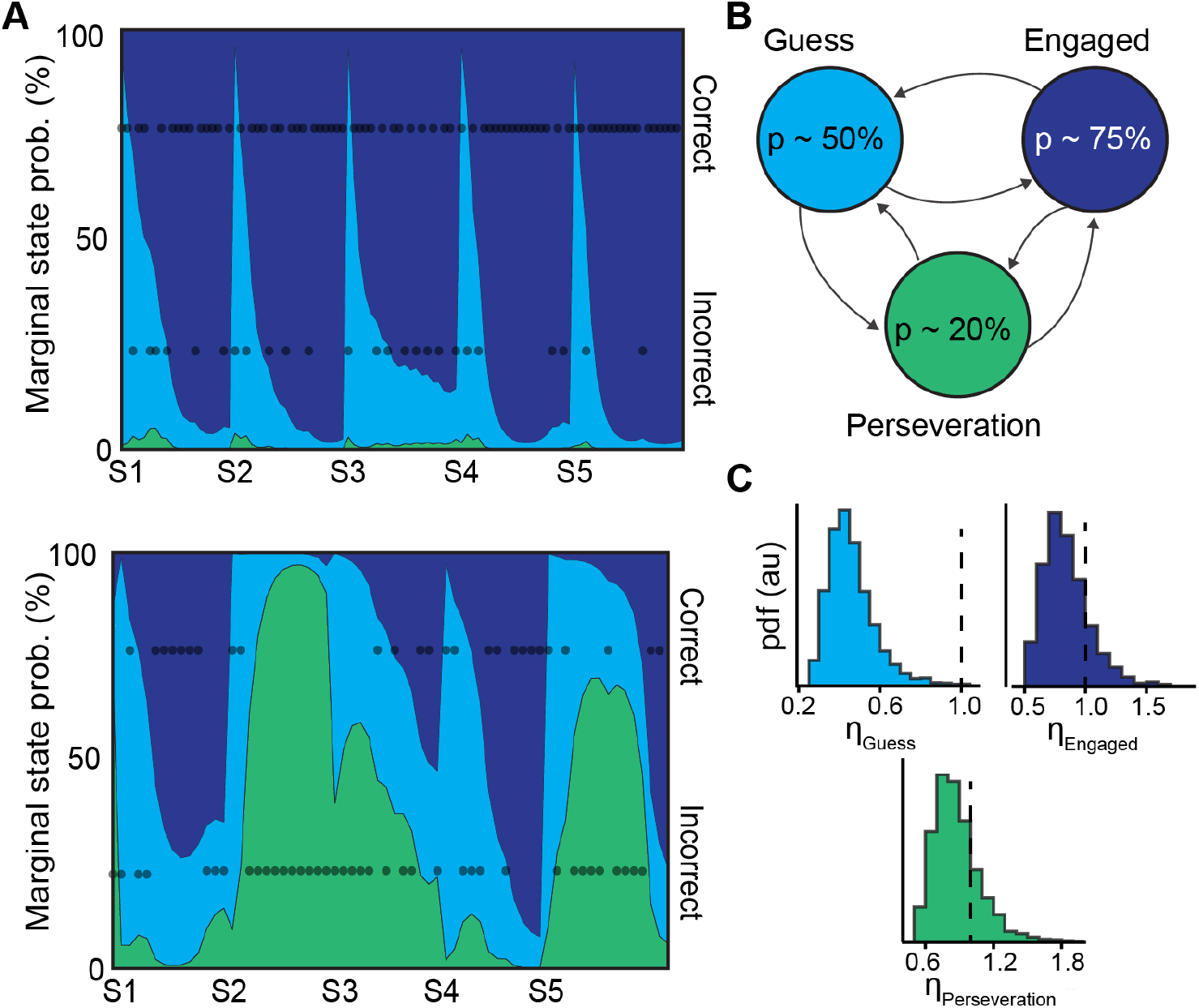
Mice switch between distinct behavioral states with differing IS rates from trial-to-trial. (A) The performance of two example animals are shown. The shaded areas represent the estimated marginal probability of being in one of three states (B) determined using a hidden Markov model, the trial-to-trial performance was partitioned into three states. The dots show the outcome (correct v.s. incorrect) for each trial. (B) The probability of a correct alternation followed a Bernoulli random variable (rounded here for simplicity of interpretation). (C) In the state labeled as “Guess,” the rate of IS in the delay zones was down-modulated by a gain term *η*_Guess_ = [0.27, 0.65], indicating a 95% HPD excluding unity. The other gain terms were consistent with unity, i.e. a neutral effect on the baseline rate.

To understand the relationship between the inferred task-related behavioral state and IS, we examined whether the rate of IS in the delay zone on each trial was different in each state. We found that the distributions of the rate of delay zone IS in each behavioral state, estimated by the Viterbi algorithm, did not share a common center location, suggesting that the rate of IS in the delay zone are related to behavioral performance (Kruskal-Wallis rank sum test *χ*^2^ approximation, p-value*<*0.0001, *χ*^2^ = 19.4, dof = 2). To estimate the magnitude of state-specific effects on IS rate, a firing rate model similar to the maze zone analysis (Fig. 4) was built to infer a “baseline” IS rate only in the delay zone for each animal, *ρ*_*a*_ (0.51 ± 0.36 Hz, n = 7 mice, mean ± 95% CI of posterior means) (Figure 6C). With the interpretation of a gain of 1 being a neutral effect, the model predicts that the “guess” state (*η*_Guess_ = [0.27, 0.65], 95% HPD credible interval, N = 504 = 7 animals × 5 sessions × M trials/day, where M is different for each animal on each day) was associated with a significant reduction in delay zone IS, while both perseveration (*η*_Perseveration_ = [0.51, 1.26]) and engagement (*η*_Engaged_ = [0.51, 1.22]) IS rates were not modulated and were thus relatively high (Figure 6). These results indicate that both engagement and perseveration are associated with baseline interictal activity during the delay period, and raises the interesting possibility that IS are mimicking delay phase SWR replay dynamics, but that the content of replay is either helpful (engagement) or harmful (perseveration). In contrast, when there is no information available, reflected by suppressed IS rates, the animal resorts to guessing.

### A simple model of interictal spikes and hippocampal place-coding reveals major differences between SWR and IS

Replays during SWR^55,56^ are thought to be important for prospective planning and consolidation of recent actions^28,32,57,58^. We sought to assess the plausibility that IS during behavior (at reward and on outer arms) interfere with mechanisms of spatial memory, especially in regards to replay events during inter-trial periods (i.e. during the delay phase). Therefore, we built an idealized model of CA3 and CA1 place coding. We modified an existing model of place coding induced by spike-timing dependent plasticity (STDP;^59–61^) to include IS which were simulated by delivering bursts of spikes to CA3 pyramidal cells (Figure 7A-C). A single burst was delivered per trial in the same relative location in the track. In the model, a mouse “explores” a linear track where it can go left or right with 90% chance of picking the opposite of the last trial (Fig. 7C) and is “teleported” back to the center of the maze to begin the next trial. After training with STDP, the spontaneous network activity was then studied to get a general sense of high frequency oscillation (HFO) dynamics in the epileptic network. Networks that received interictal-like pulses on the maze produced larger amplitude and higher frequency HFOs compared to control networks (Fig. 7D-G). Simply by including interictal-like stimuli during training, the network spontaneously generated population events that recapitulated the major qualitative differences observed in the LFPs of ripples and pathological HFOs^23^.

**Figure 7:**
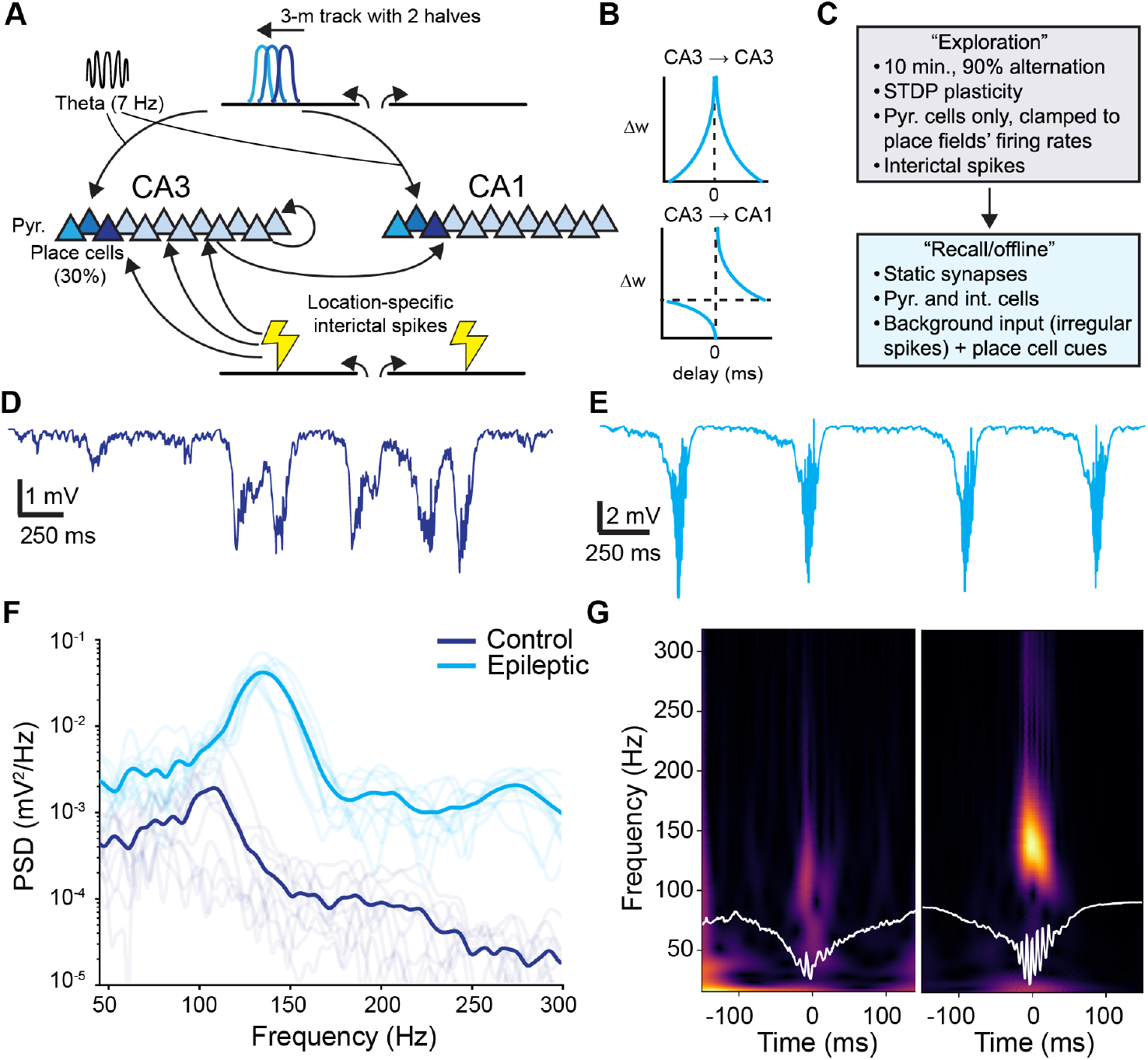
Providing IS-like input to a model of hippocampal place coding induces IS with spontaneous high-frequency oscillations (HFOs). (A-C) A schematic of the “exploration” phase of the spiking model. The virtual track was divided into two halves and interictal spikes were delivered at two locations shown schematically by the lightning bolts. (B) The STDP weight updating rules for pyramidal cell synapses during exploration are shown schematically. Representative simulated LFP traces generated from a (D) control and (E) epileptic network are shown with scale bars (250 ms and 1 or 2 mV, respectively). (F) 10 replica networks were created and the resulting spectrograms of their spontaneous replay-like bursts are shown with the group means in dark lines. (G) The mean replay oscillations from one replica network are shown along with their continuous wavelet transforms.

We then studied the spiking content of spontaneously generated replay events in the model. Like previous reports^59,60^, we observed spontaneous “replay” events of place cells in the offline state in control and epileptic networks (Supp. Fig. S4A). We used the population vector approach^62^ to reconstruct the maze positions represented by the network activity during each replay event and in each simulated subfield (Supp. Fig. S4B). Like in previous reports using similar models^59,60^, such replays were generally longer lasting and involved longer trajectories than are observed in real data, but nonetheless give a lens for comparing between control and epileptic networks.

### Spatial distribution of IS during simulated online exploration affects the quality of offline replay events

Our *in vivo* experiments showed significant variability in the distributions of IS on the maze (Fig. 3A and B). We were interested in how the spatial distribution of IS during exploration of the maze impacted the content of delay zone replays that replay remote locations (like reward locations). We simulated “cued” replay by stimulating a subset of place cells with a brief pulse of activity to induce a population event^59^. Such cued replays were performed in two cases that reflected the two extreme patterns of IS distributions we observed in our real data (Fig. 8A): the first regime (high spatial information) where simulated IS were delivered at the same relative locations on the virtual maze (like in Figures 7 and Supp. Fig. S4) and a second regime (low spatial information) where the location of each IS was varied randomly from trial to trial. We considered the location of the IS in the high information case as a ‘reference point’. Then, the relative spread of replay content beyond the cued zone was measured as 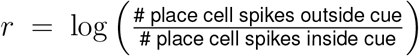 as a function of distance between the place fields corresponding to cued cells and the reference point (Fig. 8A and B). In the high spatial information case, generalization of the replay beyond the cued zone (*r* ≫ 0, i.e. beyond the case where # place cell spikes outside cue = # place cell spikes inside cue) was restricted to cue distances *<* 30 cm from the reference point. In the low spatial information case, generalization occurred at all cue distances from the reference point (excluding edge effects *>* 70 cm). In the *<* 30 cm region, the *r* values of high and low spatial information converged (Fig. 8B). These results suggest that when IS are scattered across the maze, the network is unable to generate precise replays during the delay phase of the working memory task. For example, the animal would not be able to replay previously visited reward locations (or any other locations on the maze) in isolation. Such corruption of replay could drive the low performance we observed in animals that had IS with low spatial information.

**Figure 8:**
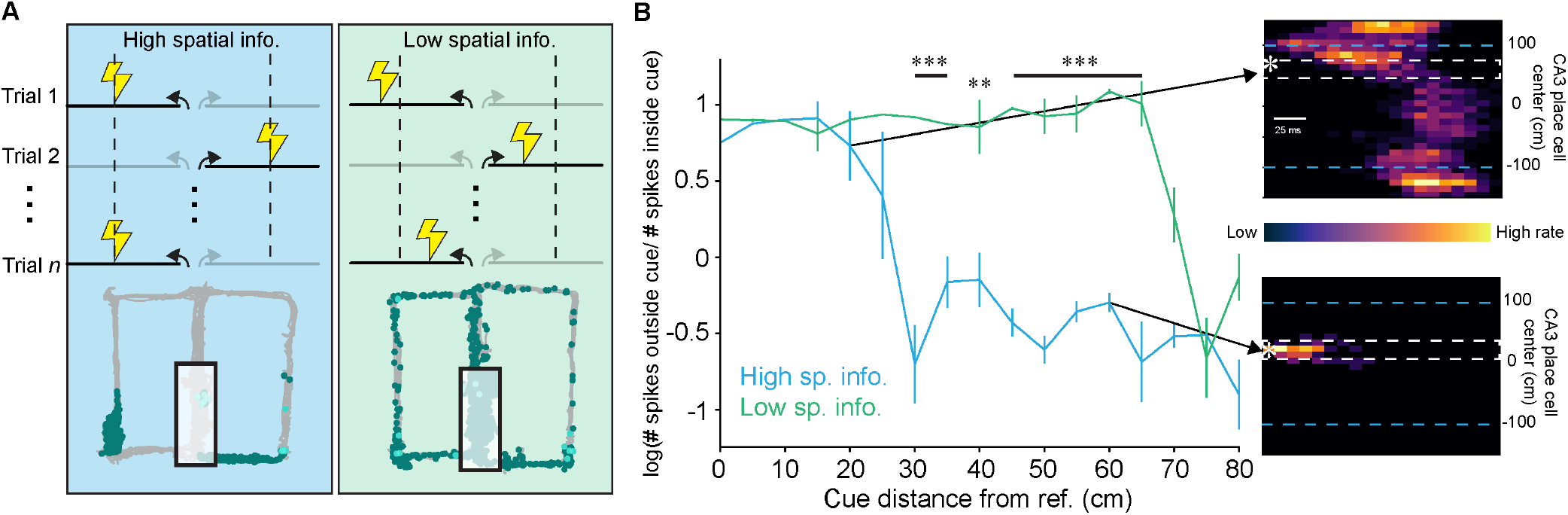
Low spatial information IS distributions lead to globally-impaired cued-replay. (A) A schematic demonstrating a simulated network with IS carrying high spatial information (left, blue) and the same for IS carrying low spatial information (right, green). The reference point is shown as a dashed box. In the low spatial information case, one IS (lightning bolt) was delivered at a random location on the maze which was changed for each trial. The mazes below show representative sessions and their IS/BIRDs (colored as in Fig. 3) with IS distributions which resemble these two statistical regimes (duplicated from Fig. 3 m1 d1, and m6 d1 left and right, respectively). The delay zone (boxed region) is not considered. (B) A cue zone was varied along one limb of the maze with respect to the rference point. The ratio of elicited place cell spikes outside and in the cue-zone was computed. The error bars show the standard error of the mean (SEM). The drop after 60 cm in the low spatial information case is likely due to edge effects from the “choice point” where simulated trials were initiated. The insets show schematic examples of cue-elicited spikes in each to extreme cueing situations. A one-way ANOVA was conducted between each pair of 10 simulations at each cue zone, ** p *<* 0.01, *** p *<* 0.001, p-values adjusted using Benjamini-Hochberg false discovery rate.

## DISCUSSION

### IS affect active encoding of spatial memory

Our recordings in freely behaving epileptic mice reveal that interictal spikes (IS) occurred frequently during a hippocampal dependent spatial working memory task (Fig. 1 and 2). IS rates were augmented during the active encoding phases of the task; both proximal to reward sites and also as animals ran down maze arms (Fig. 3). These two IS patterns had different impacts on working memory performance. When IS were spatially unrestricted, and thus carried low spatial information, animals performed poorly. Our data suggest that BIRDs which sustain themselves during locomotion (Figure 3C & D) “smear” IS across the maze and are responsible for the low spatial information. On the other hand, when IS near reward sites were distinct in LFP shape from other IS on the maze (Fig. 5), which is indicative of these IS reliably engaging ensembles with spatial information, animals performed well.

First, considering the former case where IS were spread across the maze, our observations raise the possibility that altered rates and phase-of-firing of inhibitory neurons during theta-states, like described in^63^ and^64^, allow IS to transiently break through even during locomotion. The mechanism allowing IS encroachment on theta states may be impaired inhibition in epilepsy, which may share commonalities with experimental blockade of CA2 inhibition of CA3 that enables the generation of IS-like events during theta^36^, referred to by the descriptive name of place-seizures or “pleizures”^65^. Another possibility is that cholinergic drive, which is typically high during running and is known to inhibit population synchrony^35^ might be reduced in epilepsy. Our result that the spatial information of IS was correlated with memory performance (Fig. 3B), suggests that the smearing of IS by BIRDs is detrimental to memory performance. The detrimental effect may be a consequence of aberrant activation of place cells during those IS that occur during running, which according to^23^, significantly reduced individual cells’ spatial information. We hypothesize that ensembles formed by spatially non-specific IS are a mechanism that contribute to the observed reduction in place field specificity and stability in epileptic mice reported in several studies^64,66–70^.

In the latter case we report IS being restricted to reward sites and exhibiting reward related changes, which taps into an interesting line of investigation between reward and replay-based memory mechanisms. We find a suite of reward-related changes in IS that mirror those which have been reported for SWR. This includes that IS rates are elevated in reward zones (Fig. 4), which may be an analog to reward zone related increases in SWR that have been extensively explored in healthy animals^27,31,32,71^. We also see that the amplitude of IS are larger in rewarded contexts (Fig. 5D) which could be linked to mechanisms which drive increases in number of pyramidal cell recruited to SWR in rewarded contexts in healthy animals^27^. Finally, we find the ability to discriminate IS LFP waveforms triggered in reward zones from those triggered in other locations predicts memory performance (Fig. 5B-C), which to our knowledge has no analog for SWR that has been reported. There is a connection between reward-related neuromodulation and epilepsy in general^72,73^. Dopamine and serotonin receptors are a potential target for anti-convulsant drugs^74–76^. Dopamine is also of interest to IS specifically, since in slice preparations, adding dopamine agonists has been shown to increase the rate of epileptiform bursting^77^ and its spread across cortical tissue^76,78^. Future work should focus on dissecting whether activity in the dopamine system can explain the reward-related changes in IS rate and waveform changes we have observed, and whether the spiking content underlying IS is also modulated by reward in a way that explains performance.

### IS both aid and interfere with memory-based planning during inter-trial delay phases

Our modeling results suggest that errors made in different behavioral states can be explained by the content replayed in the hippocampus during the delay period. While we do not directly observe the spiking content replayed in each IS *in vivo*, the decoding analysis in Figure 5 suggests that working memory may depend on generating delay and choice zone IS which have features that are distinct from those of reward zone IS. Furthermore, the cueing simulations in Figures 8 suggest a biologically plausible mechanism for error generation. In the case of Perseveration, spread of replay content to un-cued areas during IS in the delay zone could lead to the mouse repeatedly visiting the last visited reward area due to a failure to form a cognitive representation of state transitions needed to complete the task efficiently^79,80^. However, if the replay during an IS remains contained to the cued area (perhaps in the Engaged state, or if the animal has “high information” IS spatial distribution as like in Fig. 8) this could enable the ability to make optimal plans^58,81^ or to maintain an accurate cognitive map for alternation behavior^32^, or a mixture thereof^82,83^. In other words, the IS-induced replay could serve a role analogous to SWR-mediated replay under certain conditions but can also generate completely pathological activity depending on the patter of IS elsewhere on the maze.

### IS gain estimation framework may simplify comparisons across epilepsy models

To get a handle on behavior/location- and state-dependent changes in IS rate, we used hierarchical Bayesian models which factorized firing rate into animal- and zone- (Fig. 4) and state-dependent terms (Fig. 7). This factorization in the model’s structure relies on two key assumptions to handle variability in the IS rate: (1) that there is indeed some enduring “baseline” firing rate unique to each animal and (2) that the magnitude of modulations applied to the baseline are shared between animals. Furthermore, we relied on weakly informative priors to regularize estimation of these parameters to values that were physiologically plausible based on video-LFP monitoring data and on the range of values reported in the literature for animal models^23,37^ and those found in human epilepsy monitoring studies^84–86^. This model structure allows us to separate inter-animal variability in IS rate from potentially meaningful fluctuations around this mean value that is “universal.” Therefore, the variables of study here are not the IS rates or burdens themselves which fluctuate between distinct regimes^37^, but instead the latent unit-less modulation factors *η* in equations 8 and 17. This hierarchical paradigm may allow for more meaningful translational comparisons (i.e. between animal models of epilepsy or between species) of IS-induced memory deficits since variance introduced by subject- or systematic disease/model-differences can be accounted for as a part of the grouping structure of the model^87^. The unitless paradigm could also render meaningful estimates of *η* even when IS rates are highly variable between subjects or when IS rate is underestimated because monitoring time is limited due to clinical factors^85,88,89^.

### Limitations and future directions

Our study makes several predictions about the impact of IS on working memory. First, although we did not record single units, our data suggest several possibilities about the relationship between IS and single cell dynamics. Several studies in rodents with TLE have revealed disruption of single-cell properties including reduction in place field specificity and stability^23,64,66–70^, contamination of phase-of-firing relationships to underlying theta and gamma oscillations^63,64,70,90,91^, and aberrant post-ictal remapping^92^. An interesting possibility is that IS, especially those that encroach on theta states^93–95^, contribute to the development of such single cell pathology. In addition, with follow up single unit studies, our predictions that hippocampal replay becomes generalized when IS are unrestricted during theta states can be directly tested. It will also be important to explore whether IS show the same impacts on working memory in female mice, as this study was limited to male mice. Finally, a key finding of this study is that IS are not always negative for memory processing - and in fact, at times seem to functionally replace SWR. Such complexity indicates that future studies aimed at targeting IS to ameliorate memory deficits will need to be ‘smart’. For example, studies employing optogentic blockade of all IS, versus selective blockage of those deemed more pathological will be essential to determine the proper course of therapeutic intervention.

## Acknowledgments

This work was funded by NIH R01 1R01NS128222-01 (to L.A.E.), American Epilepsy Society Grant 835029 (to L.A.E.), VolswagenStiftung Freigeist Fellowship (to L.A.E.), and National Institute of Neurological Disorders and Stroke Grant T32 5T32NS045540-20 (to J.D.Y.). The authors thank all members of the lab for their support. We also thank Jonathan Ewell for reading the manuscript and providing feedback.

## Author contributions

Conceptualization, L.A.E, M.P., and J.D.Y.; methodology, L.A.E, M.P., G.T., and J.D.Y.; investigation, L.A.E, M.P., L.K., and J.D.Y.; writing – original draft, L.A.E and J.D.Y.; writing – review & editing, L.A.E, M.P.,L.K., G.T. and J.D.Y.; funding acquisition, L.A.E; resources, L.A.E. and M.P.; supervision, L.A.E.

## Lead contact

Requests for further information and resources should be directed to and will be fulfilled by the lead contacts, Justin D. Yi and Laura A. Ewell.

## Data and code availability

All code and data needed to reproduce the findings will be made available upon publication of this manuscript.

## Experimental model and study participant details

### Subjects

All experimental procedures were performed as approved by the Institutional Animal Care and Use Committee at the University of California, Irvine and according to National Institutes of Health and institutional guidelines or following European (2010/63/EU) and federal law (TierSchG, TierSchVersV) on animal care and use and approved by the county of North Rhine Westphalia (81-02.04.2018.A006/2 - mittel). All the experiments were performed using male C57BL/6 mice (Charles River,). All mice were single housed under a 12 h light/dark cycle, in a temperature (22 ± 2°C) and humidity (55 ± 10%). Food and water were available *ad libitum* except for during the working memory task period when mice were either food restricted to maintain 85% of their initial weight or given a 2 % citric acid water replacement when a sugar water reward was given. All efforts were made to minimize pain and reduce the number of animals used.

## Method details

### Kainate Induction of Chronic Temporal Lobe Epilepsy

Kainate injections were performed in 3-month-old C57BL/6 male mice. In one laboratory, mice were anesthetized with an intraperitoneal injection (0.1ml/10g body weight) of Ketamine (0.1 ml of 1g/ml; Bela-Pharm GmbH & Co. KG), Dormitor (0.1 ml of 1mg/ml Meditomidinhydrochloride; Orion Pharma) and Sodium chloride (0.8 ml of 0.9%; Fresenius Kabi Deutschland). Analgesia (5 mg/kg of Gabrilen, Ketoprofen) was given subcutaneously 30 mins before the surgery, and Xylocaine (AstraZeneca, Germany) was used for local anesthesia. In the other laboratory, anesthesia was induced with 3-4% isoflurane and maintained at 1-2% isoflurane. Lidocaine (2 mg/kg Patterson Veterinary Supply, USA) was used for local anesthesia. Baytril (0.5 mg, bacon flavored tablet, Bio-Serv) was used for post-operative antibiotics 5 days post-op.

Stereotactic injections were performed using a stereotactic frame (Kopf) and a microprocessor-controlled minipump (World Precision Instruments, Sarasota, Florida). 70nL of 20mM Kainate Acid (Tocris Bioscience) or saline was injected unilaterally into cortex above right hippocampus (M/L =1.5mm; A/P =1.9mm; D/V=1.1 mm from skull surface at bregma) using a 10 mL Nanofil syringe (WPI). For animals anesthetized with Ketamine, after suturing, the antagonist Antisedan (5mg/ml Atipamezolhydrochloride (Orion Pharma) was injected interperitoneally (0.1ml/10g body weight). The incision was covered with an Antibiotic-Cream, Refobacin (1mg /g Gentamicin) or Neosporin First Aid antibiotic cream. Immediately after surgery we gave 1 ml of a 5% Glucosteril solution subcutaneously. Four hours after surgery, status epilepticus was terminated using diazepam (10mg /2ml, Ratiopharm) injected subcutaneously (0.1 ml/20 g body weight), or lorazepam (7.5 mg/kg, MWI Veterinary supply) injected subcutaneously. Ketoprofen or carprofen (5 mg/kg, Rimadyl, MWI Veterinary supply) was also injected subcutaneously on the three following days to mitigate pain. Animals were left to rest for at least 1 week before starting handling.

Kainic Acid (Tocris Bioscience, ItemNo: 0222:) was prepared by combining 50mg of KA powder with a 40mM Sodium hydroxylate solution to get a stock solution of 40mM Kainate. Aliquots were stored at -20°C and mixed 1:1 with 0.9% NaCl solution to obtain 20 mM KA.

### Tetrode Recording

Double bundle microdrives (axona) comprised two bundles of 4 tetrodes separated by 3 mm to target bilateral hippocampus. The tetrodes were made of tungsten wire (Tungsten 99.95%, California Fine Wire Company) and plated with a gold solution to have impedance ∼200 kOhms. To implant the microdrives, mice were injected with the analgesic buprenorphine (0.05 mg/kg body weight) and ketoprofen (5mg/kg body weight) to reduce pain. 20 minutes later, mice were anesthetized initially with 3-4% isoflurane using an oxygen/air mixture (25/75%), placed on a regulated heating plate (TCAT-2LV, Physitemp) to retain the body temperature at 37°C, and head-fixed in a stereotactic frame. Anesthesia was performed via a mask with isoflurane 1-2% at a gas flow of about 0.5 ml/minute. After removing the skin and other tissues from the skull, a layer of Optibond (OptibondTM 3FL, KERR) was applied. Reference and ground screws were placed anterior to the bregma. Two craniotomies were drilled for tetrode implantation bilaterally (−2 mm AP, ± 1.5 mm ML) with a dental drill. After removing the dura, tetrodes were placed in 70 % ethanol for two minutes before being implanted in the cortex above the hippocampus (∼ 0.6 mm DV). After placing the tetrodes, they were covered with heated gelatinous paraffin to protect them from the dental cement. Paraffin was made with 40 g of solid wax and 50 mL oil that were mixed at 100°C. The microdrive was fixed in place using dental cement (Paladur powder and liquid, Kulzer). Mice were injected with glucose monohydrate (Glucosteril, Fresenius Kabi Deutschland; injection volume 0.25 ml, s.c.) and were kept single-housed on a heat-pad. They were carefully monitored twice daily and injected with the analgesic ketoprofen (5mg/kg) to reduce pain on the following four days. One week after implantation, LFP recordings were acquired using a Neuralynx system (Digital Lynx 4SX, Sample Rates 32 kHz, filtering 1-8000 Hz) and Cheetah 6.4.1.

Over several weeks, tetrodes were turned to the following configuration. On each side, one tetrode was positioned in the cortex for reference, complemented by three tetrodes in left and right hippocampus spanning CA1 to the dentate gyrus.

### Linear probe recording

4 weeks after Kainate injection, a high-density linear silicon probe (Neuronexus, H64LP A1×64-Edge layout, 64 channels, 20 *µ*m spacing) was implanted in the cortex above the right hippocampus (AP -1.9 mm, ML +1.6 mm, DV 0.8 mm). Anesthesia and post-operative care was done as for KA injections described above with the exception including dexamethasone (MWI Veterinary Supply, 2-4 mg/kg, i.p.) during implantation and buprenorphine (MWI Veterinary Supply, 0.05 mg/kg, s.c.) and carprofen (5 mg/kg, Rimadyl, MWI Veterinary supply) was used for peri-operative analgesia. After the mouse recovered for 1 week, the probe was lowered manually over 5 days using a microdrive (3D Neuro – R2Drive,^98^) to a depth of approximately 2.4 mm. During all recording sessions, the probe was connected to an OpenEphys (OE) Acquisition Board via a 64-channel Intan Omnetics headstage. The signal was recorded using a custom Bonsai workflow, where the OE board output was recorded using an Intan Rhd2000 Evaluation Board Node sampled at 30 kHz^99^.

### Spatial Alternation Task

Memory task training started two weeks after KA induction. The maze apparatus is a figure-8 shape (dimension 80 × 90 cm). Mice were trained to perform spatial alternation to receive sugar pellet rewards (200 mg, Test Diet). Mice were food restricted to 85 % of their baseline weight. Training consisted of three phases: (1) Habituation (2) Forced Alteration (3) Free Alternation. During habituation the mouse freely explored the maze that was covered with 9 sugar pallets (3 per arm). The Habituation phase was continued daily until the mouse ate all pellets in under 5 minutes. During the Forced Alternation phase mice were guided to alternate between right and left side of the maze using barriers placed on the maze by the experimenter. During the Free Alternation phase, the mouse was allowed to freely choose between visiting the two sides of the maze and only visits the opposite arm from the previous trial were rewarded with a sugar pellet and considered ‘correct’. Mice reached training criteria when they performed*>* 80% correct choices on 2/3 consecutive sessions of the free alteration phase. During the 3 phases of training, there was never a delay between trials. After reaching criteria, food restriction was terminated. The mice ate freely and rested for 5-7 days before microdrive implantation was performed. After surgery, mice were retrained to run with cables and again reached criteria before being passed to the memory testing phase. Testing comprised 5 days where mice ran 15-30 trials with delays of 30 seconds between trails. Before and after behavioral sessions, mice were placed in a monitoring chamber (glass bowl) where they were video-LFP monitored for at least three hours per day.

One mouse (implanted with a silicon probe) was implanted prior to any training and was run in an automated version of the Figure-8 maze (48 × 48 cm). One day prior to habituation, the mouse was placed on a 2 % citric acid water regiment^100^. Video tracking was controlled by a Bonsai workflow and maze doors and reward ports (Sanworks, Mouse Port Assembly) were operated by an Arduino micro-controller which interfaced with Bonsai. When the mouse broke an IR beam to drink, approximately 10 *µ*L of 5% sucrose water was dispensed as a reward. The training schedule was similar to that for the mice run on the non-automated maze, but involved habituation to the maze and automatic doors rather than eating food pellets.

On all behavior days, the mouse rested in a home cage immediately before and after the maze session for ∼15 minutes, during which the LFP was recorded. After the completion of all behavioral days, the mouse was video monitored in a home cage once for 14 hours overnight (6:30 pm to 7:30 am) to estimate seizure burden.

### Quantification and statistical analysis

#### Interictal spike detector

All signal processing was done in MATLAB (R2024a and R2024b, The Mathworks). Single channel LFP signals were selected based on their location being in the hippocampus (confirmed by histology) and on the amplitude of interictal spikes. The LFP was down-sampled to 1000 Hz. The sign of the signal’s skew was estimated and used to ensure interictal spikes were oriented positively regardless of the original polarity of the signal. Then, the signals were band-pass filtered (see Table 1), and peaks with a minimum prominence above a tuned threshold (Table 1) were counted as the location of interictal spikes.

#### Detector tuning

For each animal, random 3-minute segments were selected from representative 3 behavioral and 1 sleep sessions for a total of 12 minutes per animal. Windows around interictal and or ictal spikes were labeled manually using the MATLAB SignalLabeler GUI and used a “ground truth” for tuning the detector. The detector was run on this ground truth dataset and the threshold, low- and high-pass bands were varied to maximize the F_1/2_ score for each animal (Table 1). True positives (TP) were counted if the detector labeled exactly one spike within the labeled window. False positives (FPs) were either (1) any additional spikes within a labeled window or (2) any spike outside a window. False negatives (FNs) were windows containing no detected spikes. True negatives were thus not evaluated. These values were used to calculate the Precision, Recall, and F_1/2_ score using the following equations:

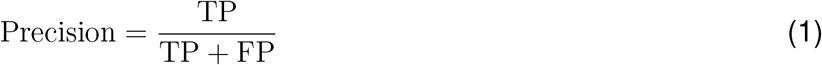

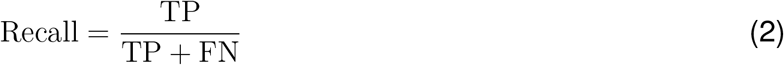

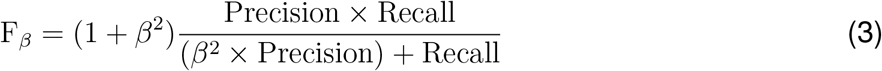

The F_1/2_ score was chosen to favor Precision roughly twice as much than Recall (*β* = 1*/*2; i.e. only selecting events that are very high above the noise floor) to avoid including non-interictal/ictal spike noise contaminating the data.

#### Binning maze zones

For a given session, the trajectory of the animal was plotted and segmented into zones. Trajectories across sessions were aligned and binned into 4×4 cm bins. Each trajectory was fitted by a rectangle and a dissecting line after calculating the coordinates of the four corners and the center of the maze. Coordinates were used to break the maze into zones with user-defined size including delay (40 cm of central arm), stem and choice (15×15 cm), outer arm, and reward zones (15×25 cm). To perform trial-wise analyses, the session was parsed into individual trials based on the sequence of entering the zones. For the automated maze, spurious “positions” that were outside the maze due to tracking errors were removed manually by inspection *post hoc*.

#### Spatial information of interictal activity

To get a sense of the “spread” of spikes on the maze, we treated the IS as if they were generated from a single “place cell” and applied spatial information analysis to its activity^39^. First, the maze was binned into a 15×15 grid, and the occupancy and number of spikes was calculated to get rates, *λ*_*i*_, and occupancy probabilities, *P* (*x*_*i*_). These were used to in the information rate (bits/s) formula provided by^39^,

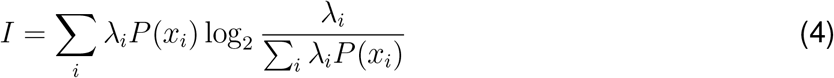

where the original integral has been replaced by a sum over occupied spatial bins, each indexed by *i*. Finally, to get information per spike (bits/spike), the quantity, *I*_spike_ = *I/*∑ _*i*_ *λ*_*i*_*P* (*x*_*i*_), was computed. This quantity was computed for each session.

To study how locomotion impacts *I*_spike_, for each BIRD, *i*, we computed the distance traveled as

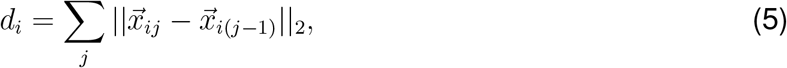

which is the sum of distances along the path defined by successive spikes indexed by *j* (compare to simple displacement from the position at the start and end of the BIRD). The operator || · || _2_ is the Euclidean norm. A generalized linear model with mixed effects (GLME) was then fit, with specification

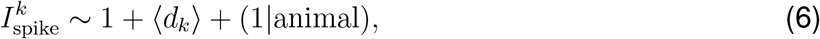

where the index *k* is a per-session index, and ⟨*d*⟩_*k*_ is the mean BIRD distance traveled within that session. The model was fitted in MATLAB using the `fitglme()` function and Gamma distribution with reciprocal link. Gamma regression was selected since 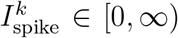 and diverged from a normal distribution when inspected on a quantile-quantile plot. For each session, the alternation task performance was also fit to compare the spatial information to performance as

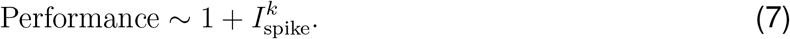

The MATLAB `fitglm()` function was used to fit the regression and dispersion was estimated from the data.

#### Zone-specific IS Rate analysis

Mouse location was binned into zones of the maze: “Delay,” “Choice,” “Reward,” and “Outer Arm” regions of interest. For each zone, the observed spike counts were calculated by calculating the number of IS in a given zone. These observed counts were compared to Expected counts which were calculated by multiplying the % of time in each zone by the total spike count. Observed and expected spike counts were compared with a *χ*^2^ test.

To estimate the zone-specific influence on the observed IS number of spikes for a given animal in each zone, *S*_*z,a*_, we employed a Bayesian approach to infer zone-specific “gains,” *η*_*z*_, which were applied to an animal-specific “baseline” IS rate, *ρ*_*a*_ as:

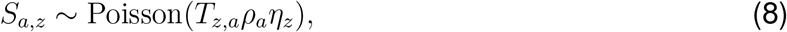

where *T*_*z,a*_ is the number of seconds in the zone *z* spent by animal *a*. (Note the correspondence of this parametrization to that of a standard generalized linear model (GLM) with a Poisson distribution and log link function via the identity, 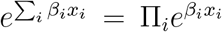, where *β*_*i*_ and *x*_*i*_ are generic regression coefficients and predictors.) The parameters to be estimated had priors of the following form:

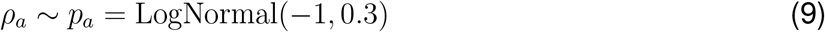

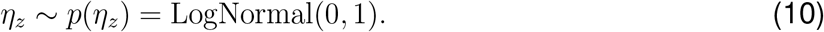

Therefore, the posterior distribution was expressed as

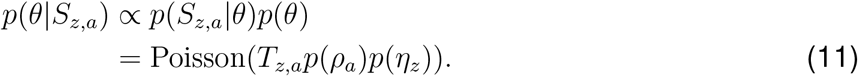

The model was specified in the probabilistic programming language Turing.jl in Julia (version 1.10.2,^101^), with packages managed with DrWatson.jl^102^. Four independent chains each run for 1000 iterations with 500 warm-up samples were run using the No-U Turns Sampler (NUTS,^103^) with a target acceptance ratio of 65% to estimate a posterior distribution for the parameters. 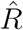 values and effective sample sizes (ESS) were checked to ensure convergence, mixing, and sampling efficacy of the Monte Carlo Markov chains.

95% credible intervals (1-*α*) were estimated for each parameter by using the highest posterior density (HPD) method. The credible interval for each *η*_*z*_ was compared to a “null” value of 1, and for those which did not overlap with 1, a “significance level” was estimated by lowering the (HPD) threshold *α* until the credible interval contained 1.

To assess the model fit, samples from the posterior predictive distribution were taken and used to generate “replicates” of the data, 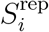.^104^. The distribution of means of the replicates were compared to each observed data point for agreement. As a check of model specification sensitivity, mean squared errors were calculated for this model using the replicates and observed data, and then compared against a “clamped” model fit where *η*_*z*_ = 1 for all zones. The two models had Akaike information criterion (AIC) values of approximately 1.30 × 10^4^ for the full model versus 1.48 × 10^4^ for the clamped model. Thus, the full model was retained for its interpretability and improved prediction performance.

#### IS LFP embedding and classification

For each IS that occurred on the maze, the single channel LFP signal was extracted ± 100 ms from the detection time. Then, the LFP was down-sampled to 2000 Hz and transformed to a z-score. The LFPs for each animal on all delayed alternation behavior sessions were then non-linearly embedded with t-SNE (with default parameters) to get a 2-D feature vector. An bagging ensemble of trees^49,50^ was fit using MATLAB with 5-fold cross-validation (including stratification into groups with similar proportions of each discrete class) to classify whether or not the IS occurred in either reward zone based on the feature vector’s position in 2-D space. The area-under-the-curve (AUC) of the receiver operating characteristic (ROC) curve was computed and compared to the animals mean performance over 5 sessions using standard linear regression. Note that the qualitative results did not change when the classifier was trained on the full-dimensional LFP waveforms instead of the t-SNE embedding, suggesting the embedding faithfully reduces the dimensionality by persevering relevant features.

To compare the amplitudes of the IS events under different conditions, the root-mean-squared (rms) amplitude was computed for each IS. To compare across animals, the raw rms values were divided by the standard deviation of the rms for all the IS of a given animal. Two-sample t-tests were used to compare the distributions of amplitudes between IS inside v.s. outside reward zones, and IS at reward during correct v.s. incorrect trials.

#### Inferring trial-to-trial behavioral state from task performance

The efficacy of decision-making depends in part on the underlying behavioral state of the animal—whether the animal is engaged with the task or has a lapse in performance. This dependency of task performance and neural dynamics on a latent behavioral state has been modeled using models that capture auto-regressive dependencies across trials^51,105–107^.

Borrowing ideas from^51^, we modeled the trial-to-trial performance using a hidden Markov model (HMM) with states inferred from the data as follows. Consider discrete states indexed as *s* ∈{1, 2, · · ·, *N*}. The probability of an animal making a “correct” choice on trial *i* depends on the state as

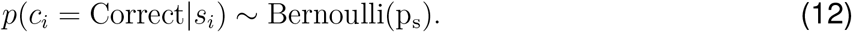

In other words, the performance is like flipping a biased coin with probability of “heads” *p*_*s*_. The value of *c*_*i*_ is considered as the “emission” of the hidden Markov chain. The state can change from trial to trial, and thus the probabilities of transitioning between different states are expressed as

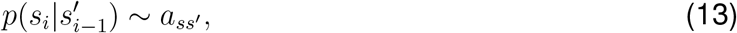

Where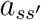 ; is an entry in the transition matrix *A*∈ ℝ^*N* × *N*^. The initial state on the first trial is drawn from

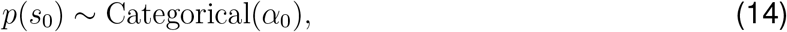

where *α*_0_ ∈ℝ^*N*^ is the probability of initializing in each of the N states. These parameters were initialized as:

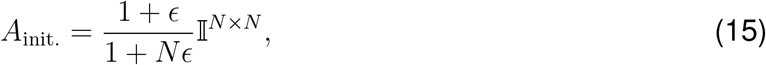

where 𝕀^*N* × *N*^ is the identity matrix, *ϵ* = 0.5 is a parameter to control the relative strength of transitions between states versus persisting within the same state, the initial state as:

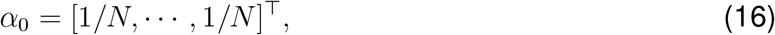

and finally, each *p*_*s*_ took one of *N* uniformly spaced values from 0.1 to 0.9.

To train the HMM, all the choice data for each trial from each epileptic animal was concatenated into a vector and the end of each session was noted. Then, the Baum-Welch expectation-maximization procedure was applied to this concatenated vector (re-initializing when a session ended) to find the optimal values of the initial state distribution, the transition matrix, and the emission probabilities for each state^108^. Using the optimized HMM parameters, the most likely state sequence given the observed choice data was computed using the Viterbi algorithm. Marginal probabilities of each state were also estimated using the forward-backwards scheme. HMM algorithms were used from the HiddenMarkovModels.jl software package in Julia^109^. This procedure was conducted for *N* = 2 and *N* = 3. The two HMMs had similar log-likelihoods after Baum-Welch estimation (−324.8 and -323.5 respectively), and so only the *N* = 3 case was retained for further analysis. Finally, the hierarchical bootstrap method^97^ was applied to estimate delay period exit times by stratifying the data into 3 states, then sampling with replacement a single trial from an animal weighted by the number of trials that animal had within that state until the a sample of the same size as the original data in each state was generated. The mean of these samples was computed for 1000 replicas.

#### Inferring behavioral state-dependent IS activity in the delay zones

The inferred marginal probabilities of each state sequence from the forward-backward algorithm,*p*(*s*_*i*_), were used as a prior to parameterize a variant of the firing rate model. The model likelihood was specified as

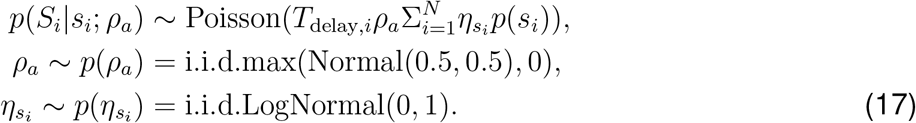

The variable *S*_*i*_ is the number of spikes in the delay period on trial i. The term 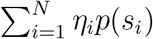 is the sum of gain terms 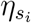 each weighted by the marginal probability of being in state *s*_*i*∼_ *p*(*s*_*i*_). *T*_delay,*i*_ was defined the time in seconds the animal spent in the delay zone at the start of the trial *i*. This model can be interpreted as applying a state-specific scalar gain 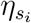 to an underlying animal-specific firing rate *ρ*_*a*_. The prior for *ρ*_*a*_. was chosen to be weakly informative of the fact that the previous Bayesian model returned animal-specific mean rates centered around 0.5 Hz, truncated at zero to exclude negative rates. Modifying the standard deviation of this prior from 0.5 to [0.1, 0.8] had no effect on the qualitative conclusions of the inferences. The model was again estimated in Turing.jl using 5000 samples from the NUTS sampler, initialized as before.

To validate the model estimated gains 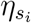, the discrete state-sequence from the Viterbi algorithm was used to group spike counts into *N* distributions. A Kruskal-Wallis test was used to compare the spike count distributions. As posterior predictive checks, the means of the posterior mean *ρ*_*a*_ values were compared to the observed mean rate of IS from the data. Also, the distribution of predicted marginal mean spike counts 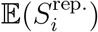 and the distribution of marginal means counts conditioned on the Viterbi-estimated state, 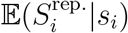, were compared to their point-estimates from the observed data, 𝔼(*S*_*i*_) and 𝔼(*S*_*i*_|*s*_*i*_). Note that 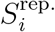 denotes samples from the posterior predictive distribution^104^.

### *In silico* model of IS and place-coding replay

#### Spiking model of hippocampal replay

To isolate the effects of IS on hippocampal coding required for behavioral navigation, we modified the spiking neural network model of place cell replay in CA3 described by^59^ and then updated by^60^ to include CA1. Pyramidal cells (pyr., n = 1250 region) and interneurons (int., n = 100 per region) in CA1 and CA3 were modeled using the adaptive exponential leaky integrate-and-fire (AELIF,^110^) model,

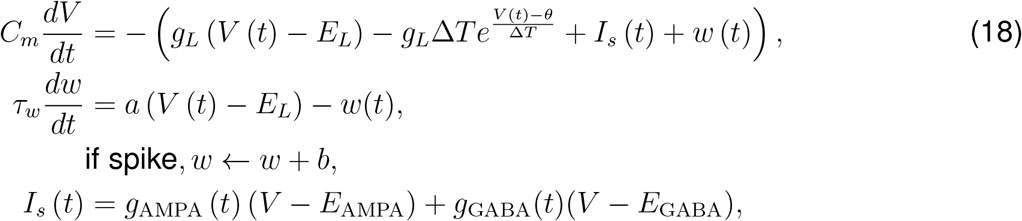

where *V*(*t*) is the membrane potential, *w*(*t*) is an adaptation current, and *I*_*s*_(*t*) are the sum of synaptic currents. Furthermore, *C*_*m*_ is the membrane potential, *g*_*L*_ and *E*_*L*_ are the leak current conductance and reversal potential, *θ* is the spike threshold, Δ*T* is the threshold sharpness, *τ*_*w*_ is the time constant for adaptation, *a* and *b* are parameters specifying how the adaptation current evolves between and following neuron spikes, respectively (see Table 9 for values which approximately correspond to those in^59^). The synaptic conductances *g*_AMPA_ and *g*_GABA_ were bi-exponential functions as in^59^, with *E*_AMPA_ = 0 mV and *E*_GABA_ = −90 mV. All neural simulations were specified and run in NEST v3.7^111^ with Python 3.12.3.

**Table 9:** Spiking neuron parameters for model equations 18.

| Parameter (units) | Value (pyr., int.) |
| --- | --- |
| $C_m$ (pF) | 180, 118 |
| $g_L$ (nS) | 4.3, 7.5 |
| $E_L$ (mV) | -75, -74 |
| $\theta$ (mV) | -24, -57.7 |
| $\Delta T$ (mV) | 4.23, 4.6 |
| $V_{\text{peak}}$ (mV) (when a “spike” is detected) | -3.25, -34.78 |
| $V_{\text{reset}}$ (mV) (reset voltage after spike) | -29.7, -65 |
| $t_{\text{ref.}}$ (ms) (refractory period) | 5.9, 1 |
| $\tau_w$ (ms) | 83.4, 178.58 |
| $a$ (nS) | -0.27, 3.05 |
| $b$ (pA) | 206.84, 0.91 |

To simulate the plasticity induced by repeated exploration of a maze environment, we adopted a modified the place- and theta-modulated spike-timing-dependent plasticity (STDP) paradigm introduced in^59^ and^60^; a similar form model was experimentally validated in^61^. The original model only considered a single 3 m long linear track, whereas our task involves alternating across two separate arms of a maze. Therefore, to understand whether the two arms are re-activated during replay separately, we modified the “exploration” paradigm to take place on two 150 cm arms pointed left and right, with the mouse starting at the midpoint and “teleporting” back to the midpoint once it reached either end. For 10 minutes, leftward and rightward trajectories were chosen at random according to a 90% chance of alternation. The simulated mouse ran at 35 cm/s with a theta oscillation frequency of *f*_*θ*_ = 7 Hz. In both CA1 and CA3, 30% of pyramidal cells were selected as place cells and given each a place field center *x*_*i*_ drawn uniformly from the total length of the maze. During the “exploration” phase, only pyramidal cells were simulated as inhomogeneous Poisson processes with firing rates as

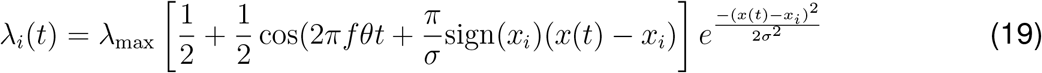

where the maximum firing rate at the center of the place field was *λ*_max_ = 20 Hz, and the width of the place field was *σ* = 7 cm. This equation encapsulates place tuning and theta phase precession^59,60^. Non-place cells fired with a mean rate of 0.1 Hz. To simulate ictal spikes which occur generally in the same location on the maze during exploration (for example, the reward ports), all CA3 pyramidal cells received a pulse of spikes at *λ*_max_ = 2000 Hz described according to

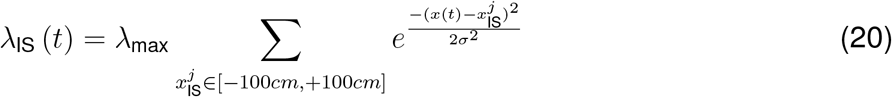

where *σ*_IS_ = 4 cm. A single spike train was drawn from *λ*_IS_ (*t*) and broad-casted to all the CA3 pyramidal cells, but each neuron only received each spike with independent probability of 1%. Since during exploration the pyramidal cell spiking was clamped to the rates above induce sequences encoded in the weight matrix tuned by STDP^59–61^ both the internal dynamics of pyramidal cells and interneuron dynamics and synapses were neglected. The normalized synaptic weights, *w* ∈ [0, 1], were updated according to standard STDP rules^112^ as

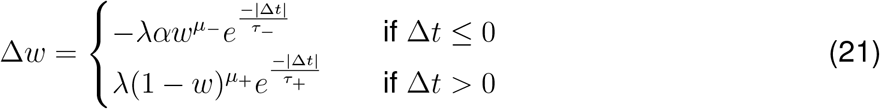

where *µ*_±_ = 0 (weight-independent updating rule) and Δ*t* = *t*_*post*_ − *t*_*pre*_, *λ* is the step size parameter, *α* is an asymmetric parameter controlling synaptic depression, and *τ*_+_ and *τ*_−_ are the time scales of facilitation and depression respectively. For CA3-to-CA3 pyramidal neuron synapses STDP was symmetrically facilitating as in^60^, thus, 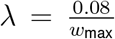, *α* = −1, *τ*_±_ = 62.5 ms, and *w*_max_ = 40 nS. For CA3-to-CA1 synapses, 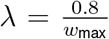 nS, *α* = 0.4, *τ*_+_ = 20 ms, *τ*_−_ = 40 ms, and *w*_max_ = 40 nS. Weights were initialized as 0.3LogNormal(0,1) nS for CA3 and 0.7LogNormal(0,1) nS for CA1. Synapses were formed between CA3 pyramidal neurons recurrently and fed forward to CA1 neurons with 10% probability for each pair of cells. The training procedure above was repeated 10 times for control and epileptic conditions to generate different replicas.

To simulate spontaneous replay during “offline” states such as the delay period between trials, the full network with pyramidal cells and interneurons with AELIF dynamics was constructed. The final pyramidal-to-pyramidal cells weights learned by STDP after all exploration trials were used to parameterize static synapses. CA3 pyramidal cells were stimulated by background activity from the dentate gyrus that was assumed to have a pooled rate of 12 Hz and synaptic weight of 20 nS. The connections between all other cell types are detailed in Table 10.

**Table 10:** Synaptic parameters during “offline” state modeled after^59^.

| Pre, Post | Weight (nS) | Delay (ms) | Rise time (ms) | Decay time (ms) | Prob. connection |
| --- | --- | --- | --- | --- | --- |
| CA3 pyr., CA3 pyr. | STDP | 1 | 1.0 | 9.0 | 0.1 |
| CA3 pyr., CA1 pyr. | STDP | 1 | 1.0 | 9.0 | 0.1 |
| CA3 pyr., CA3 int. | 0.85 | 1 | 1.0 | 9.0 | 0.1 |
| CA1 pyr., CA1 int. | 0.85 | 1 | 1.0 | 9.0 | 0.1 |
| CA3 int., CA3 pyr. | 0.65 | 1 | 0.3 | 3.0 | 0.25 |
| CA1 int., CA1 pyr. | 0.65 | 1 | 0.3 | 3.0 | 0.25 |
| CA3 int., CA3 int. | 5 | 1 | 0.3 | 3.0 | 0.25 |
| CA1 int., CA1 int. | 5 | 1 | 0.3 | 3.0 | 0.25 |
| DG, CA3 pyr. | 20 | 1 | 0.65 | 5.4 | 0.25 |

For comparing how the IS distributions on the maze affected network cueing, we simulated a case with high spatial information (the default described above) and low spatial information case by varying the location of IS uniformly over the interval [± 0 cm, ± 150 cm]. To “cue” replay, the weight of background activity was reduced to 10 nS, and CA3 place cells associated with different zones were stimulated with a 20 ms burst of spikes sampled from a Poisson process at 30 Hz with a synaptic weight of 500 nS from the simulated dentate gyrus. Cues were given to CA3 place cells on 20 cm wide intervals centered on equally spaced (5 cm) locations between 0 and 80 cm away from a reference point. The weight of each cue synapse was set to 80 nS. Only one network for each case was used, and 10 random seeds were used to initialize simulations at each cue center. Using the Scipy library, a one-way ANOVA was conducted at each cue center and p-values were corrected with a Benjamini-Hochberg false discovery rate procedure.

### Analysis of simulated LFP

The “LFP” proxy of the network was computed as the sum of all synaptic currents delivered to a random subset of 200 CA1 pyramidal cells,

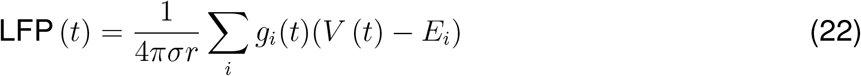

sampled at 1000 Hz where the extracellular conductivity *σ* = 0.3 S/m and the distance from each current source to the electrode was set to be *r* = 5*µ*m. The choice of these parameters only affects a scalar gain^59^. Replay events were detected by detecting peaks with a prominence of 1 mV and minimum distance of 200 samples on the lowpass filtered LFP at 200 Hz with a 7th order Butterworth filter. Once the peaks were found, 150 ms on either side of the peak were selected and used for further analysis. The power spectral density (PSD) of the LFP was estimated using Welch’s method with 256 samples per segment, 32 sample overlap, and 1024 FFT points. The PSD was computed on 5 s long segments of spontaneous activity from each of the 10 replica networks. The continuous wavelet transform (CWT) was computed on the averaged replay LFP using a complex Morlet wavelet with bandwidth of 1.5 and center frequency of 1.0 at 200 logarithmically spaced frequency bands between 10^1.2^ to 10^2.5^ Hz. To compute the CWT, only one replica network was selected and the replays over 5 s were used for averaging. Signal processing was done using Scipy^113^ and the PyWavelets package^114^,

### Analysis of simulated replay events

After replays were detected by the LFP, the place cell activity was extracted and used to reconstruct the maze position using the population vector method^62^

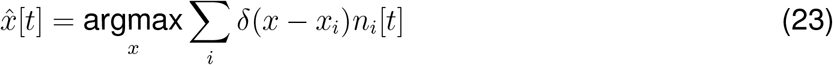

where δ (*x*) is the Dirac delta function, *x*_*i*_ is the place field center, and *n*_*i*_[*t*] indicates the number of spikes neuron *i* fired within a discrete 25 ms time bin.

## A Supplement

**Table A.1:** LFP discriminability and SI independently explain animals’ performance. Adjusted *R*^2^ = 0.86, d.f. = 4. Full model v.s. constant model *F* = 20.1, *p* = 0.008.

| Parameter | Estimate | SE | tStat | p-value |
| --- | --- | --- | --- | --- |
| Intercept | -24.6 | 14.9 | -1.65 | 0.17 |
| $\langle \text{AUC} \rangle$ | 117 | 23.0 | 5.10 | 0.007 |
| $\langle \text{SI} \rangle$ | 7.7 | 3.11 | 5.10 | 0.04 |

**Figure A.1:**
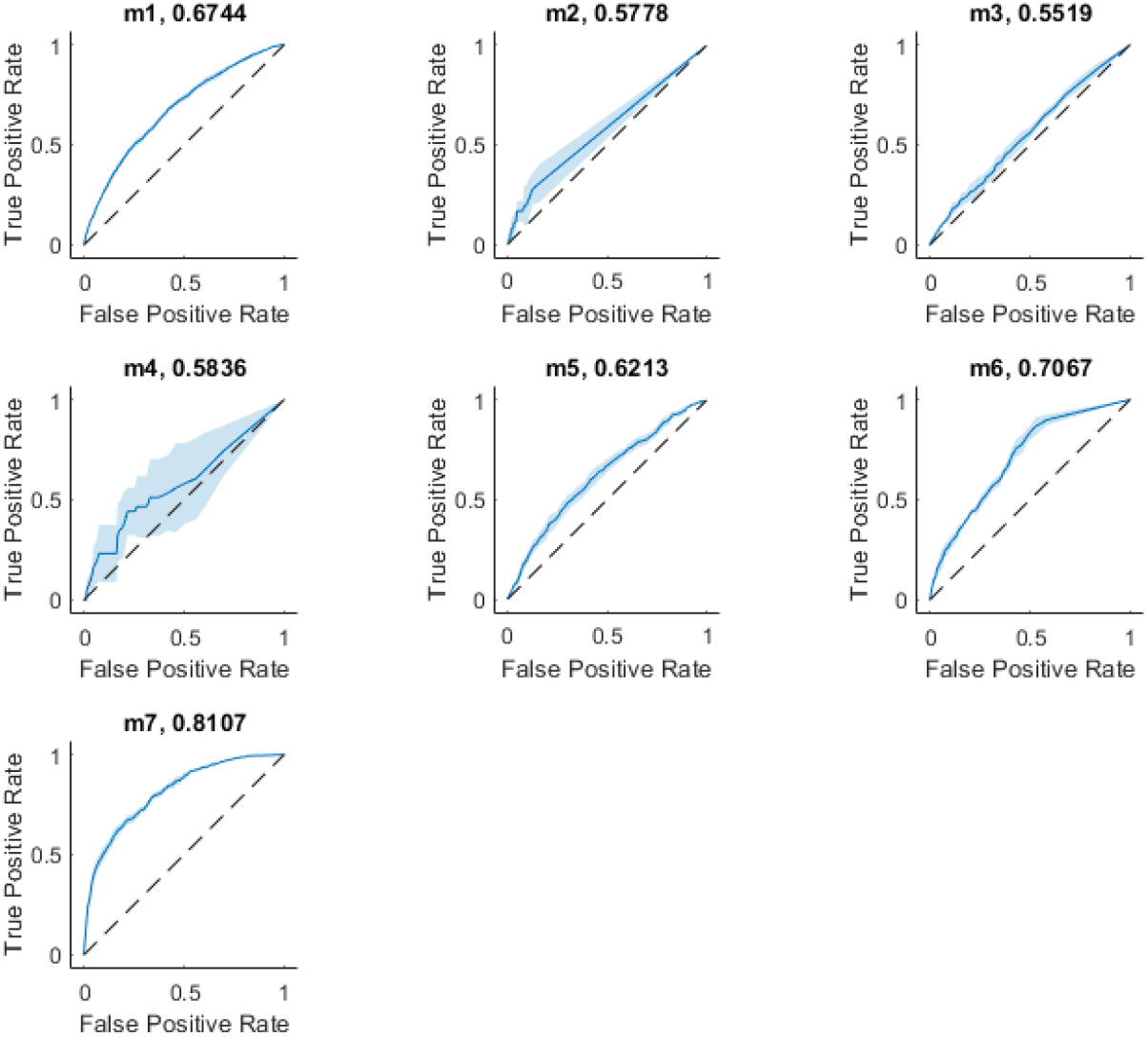
ROC curves for classifier IS performance.

**Figure A.2:**
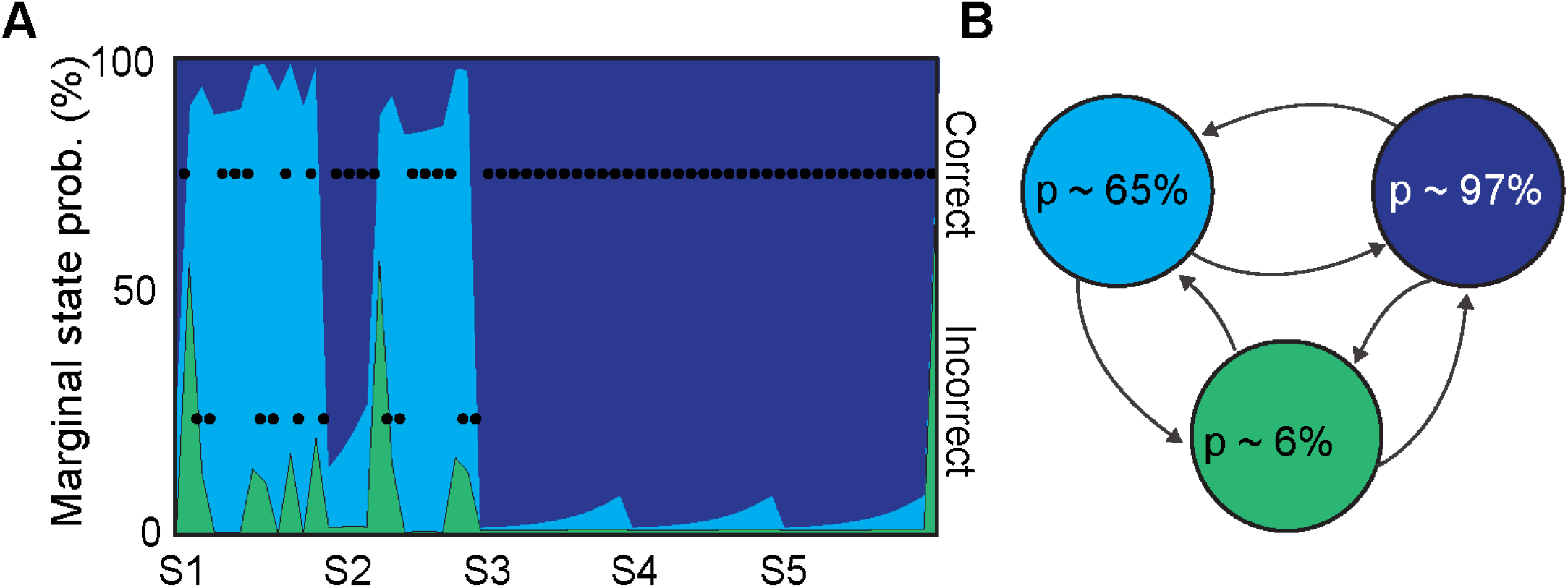
Three-state HMM fit to control animals reflects improved performance both in (A) the high proportion of time in the engaged state in this representative animal and (B) the higher Bernoulli probability of successful alternation in the engaged state than that of controls.

**Figure A.3:**
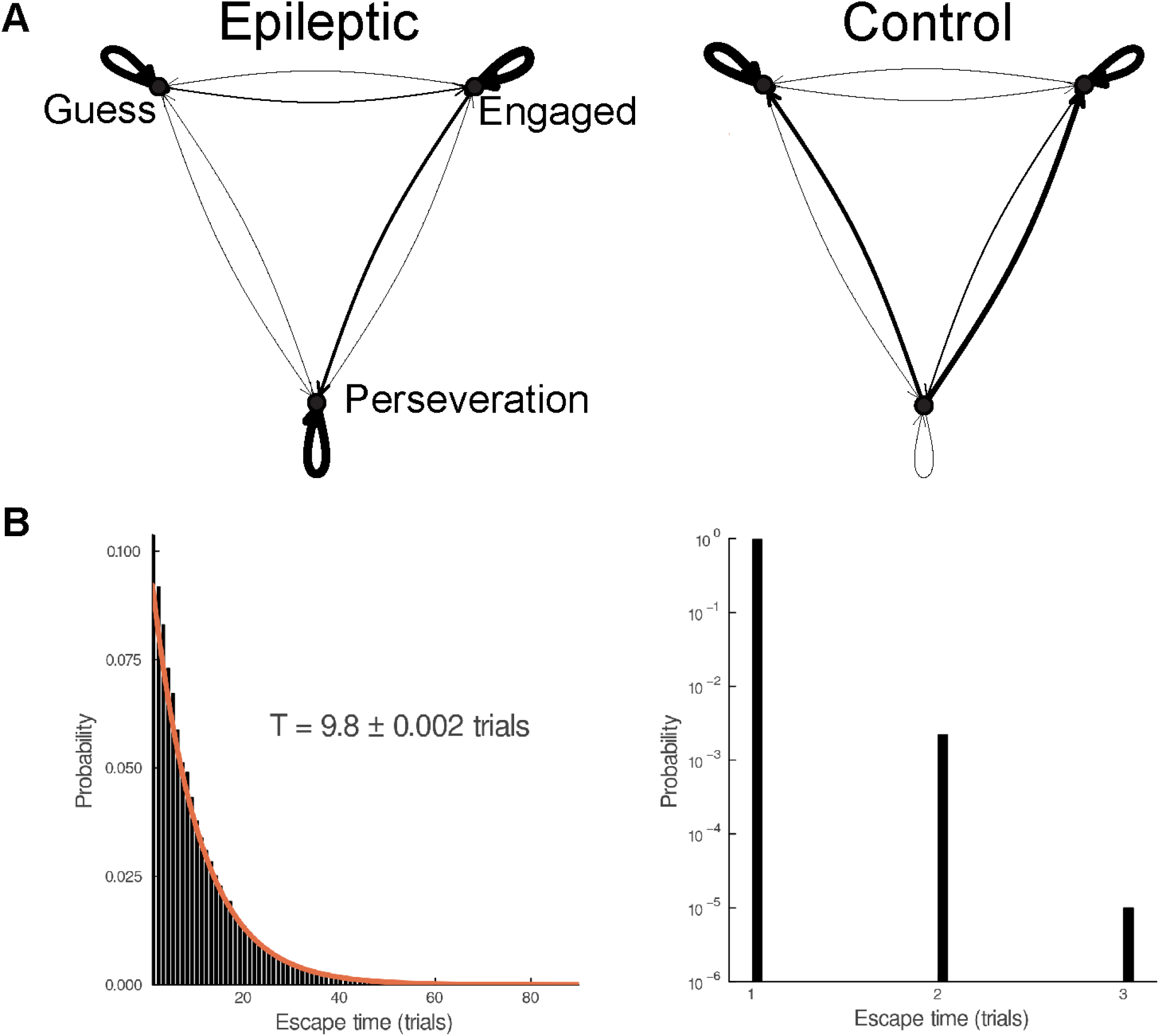
The behavioral state HMM transition matricies from epileptic and control animals. (A) The width of the arrows show the relative probability of transitioning between each state. (B) The mean escape time from the perseveration state was compared and found to be elevated in the epileptic mice, which is consistent with the magnitudes of self-transition probabilities in (A). The red line indicates an exponential fit with mean *T*.

**Figure A.4:**
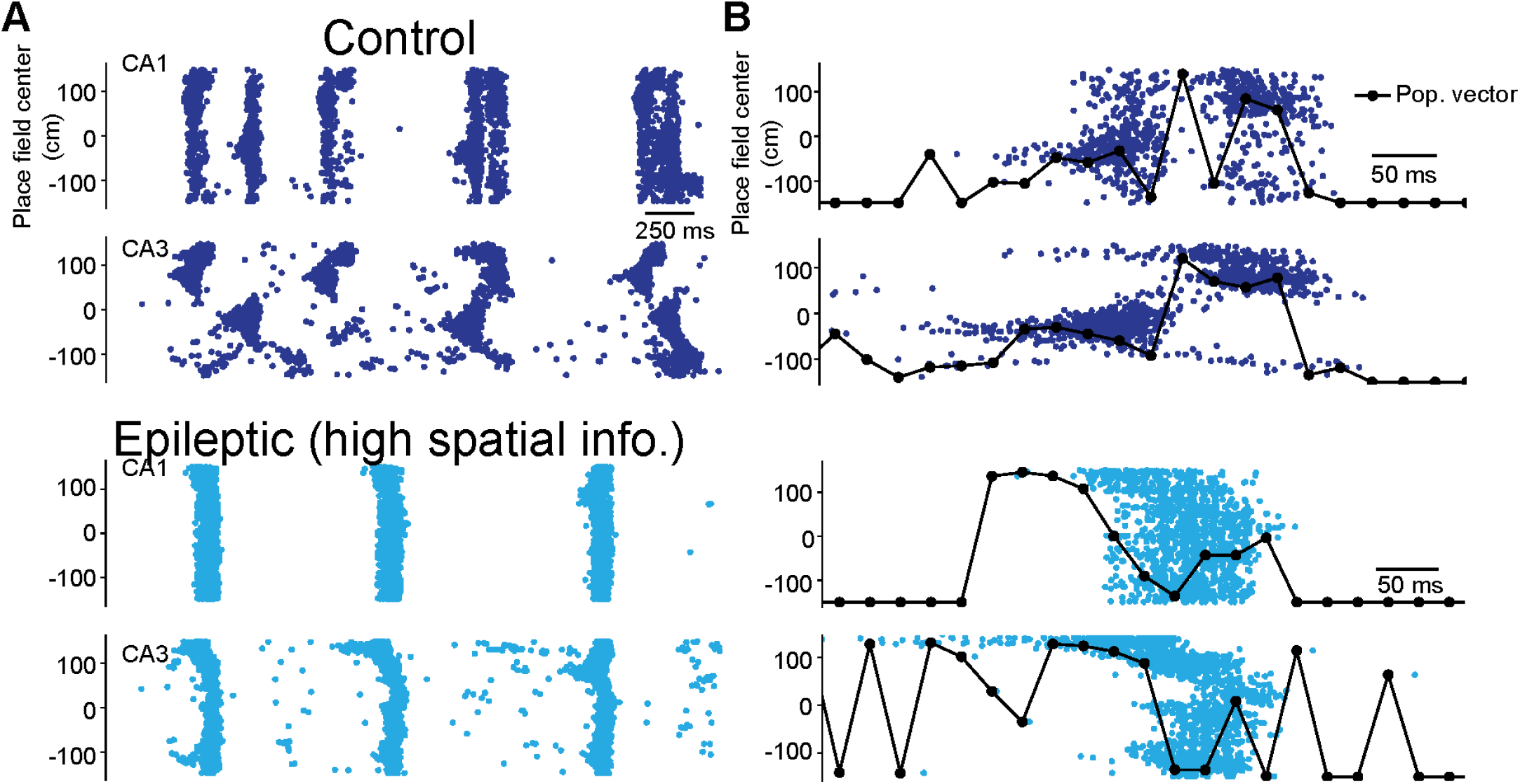
Replay events *in silico* are corrupted by recruiting off-target ensembles. (A) Raster plots of spontaneous place cell activity in CA3 and CA1 from control (dark blue) and epileptic (light blue) networks. (B) Zoomed-in single replay events, along with their respective population vector reconstructions as a black line, are also shown.

